# Nonsense-mediated mRNA Decay Inhibition Triggers Immunogenicity by HLA Class I Immunopeptide Remodeling in Non-Small Cell Lung Cancer Cells

**DOI:** 10.64898/2026.11.20.768099

**Authors:** Alexandra Emanuela Burger, Anne-Christine Uldry, Sophie Braga Lagache, Maxime Clément Boy, Jan Paul Müller, Oliver Mühlemann, Manfred Heller

## Abstract

Nonsense-mediated mRNA decay (NMD) is a post-transcriptional surveillance pathway that degrades aberrant transcripts harboring premature termination codons. Thereby, NMD also limits the abundance of transcripts encoding potentially immunogenic peptides. Here, we investigated whether pharmacological inhibition of the NMD kinase SMG1 remodels the human leukocyte antigen class I (HLA-I) immunopeptidome and enhances immune recognition of H838 non-small cell lung cancer (NSCLC) cells. Treatment with a SMG1 inhibitor (SMG1i) effectively perturbed NMD, as reflected by increased abundance of NMD-sensitive transcripts. To identify NMD inhibition-associated HLA-I peptides beyond the canonical proteome, long-read RNA sequencing and ribosome profiling were integrated to generate a cell-specific sequence database for immunopeptidomic analysis. Identified peptides were further evaluated using an extended metascore incorporating predicted HLA binding and proteasomal processing to define a high-confidence HLA-I peptide repertoire. Of the 2,728 peptides passing the 1% false discovery rate threshold, 48 were detected exclusively in SMG1i-treated samples. Of these, 32 fulfilled additional criteria for recurrence, peptide length, and predicted binding to the genotyped H838 HLA-I alleles. The observed changes in HLA-I-bound immunopeptides were not accompanied by an increase in global surface HLA-I abundance, suggesting remodeling of peptide composition rather than increased HLA-I presentation capacity. Exploratory allogeneic co-culture experiments showed SMG1i-induced IFN-γ response in partially HLA-matched CD8⁺ T-cells compared with an unmatched donor and increased expression of effector-associated genes. Together, these findings establish a cell-specific, proteogenomic, and HLA-informed framework for identifying SMG1i-associated HLA-I peptides in an NSCLC model and prioritizing candidates for peptide-specific functional validation.

## Introduction

Immune checkpoint inhibitors targeting programmed cell death protein 1 (PD-1) or programmed death-ligand 1 (PD-L1) have transformed the treatment of non-small cell lung cancer (NSCLC), providing durable clinical benefit in a subset of patients^1–3^. However, responses remain heterogeneous, and primary or acquired resistance is common^4–8^. CD8⁺ T-cell-mediated antitumor immunity relies on tumor cells presenting a sufficient repertoire of tumor-specific or tumor-associated peptides on human leukocyte antigen class I (HLA-I)^9,10^. Tumor mutational burden (TMB) and microsatellite instability (MSI) can increase the generation of potentially immunogenic neoantigens and, in selected tumor types and clinical settings, are associated with response to immune checkpoint blockade^11,12^. However, a high mutational load does not guarantee HLA-I presentation of the resulting mutant peptides. Transcript abundance and translational efficiency alone also do not reliably predict presentation. Only a limited, non-random subset of translated products ultimately generates peptides that are processed, bound HLA-I molecules, and presented at the cell surface^13,14^. Post-transcriptional surveillance and HLA-I genotype further constrain this repertoire^4,15^.

Nonsense-mediated mRNA decay (NMD) is a conserved post-transcriptional RNA surveillance mechanism that degrades transcripts harboring premature termination codons, including transcripts encoding frameshift- and splice-derived neoantigenic peptides^16^. Beyond this classical quality-control role, NMD regulates a broad range of transcript classes, thereby influencing gene expression and cellular stress responses^17,18^. In tumor cells, NMD degrades most neoantigen-encoding transcripts, thereby actively limiting the immunopeptidomic diversity available for immune recognition^19–22^. NMD inhibition therefore emerged as a strategy to expand the tumor immunopeptidome by stabilizing transcripts that would otherwise be degraded, thereby increasing the availability of canonical and noncanonical antigenic peptides^16,23–25^.

Inhibiting SMG1, a kinase essential for NMD activity in metazoan cells, has the potential to increase tumor immunogenicity. SMG1 inhibition may unmask NMD-suppressed mutation-derived neoantigens as well as peptides arising from non-mutational premature termination codon (PTC)-containing transcripts. While mutation-derived peptides may constitute tumor-specific neoantigens, non-mutational peptides are not inherently tumor-specific and therefore require assessment of whether their presentation is absent in non-malignant cells. SMG1 is a phosphoinositide-3-kinase-related kinase (PIKK) that phosphorylates RNA-bound UPF1 following aberrant translation termination^26,27^. Phosphorylated UPF1 recruits downstream NMD effectors that promote transcript degradation^26,28^. Molecular screening identified pyrimidine derivatives as potent and selective SMG1 inhibitors^29^. Subsequent biochemical studies confirmed that SMG1 inhibitors reduce SMG1-dependent UPF1 phosphorylation in vitro^30^. Cook and colleagues subsequently developed KVS0001, an *in vivo*-compatible SMG1 inhibitor derived from SMG1i-11 (11j), a compound from the same pyrimidine SMG1 inhibitor series as 11e, used in this study. This inhibitor showed increased HLA-I-associated presentation of peptides derived from NMD-regulated proteins on human cancer cell surfaces^25^. Vendramin and colleagues further showed that SMG1 inhibition reshapes the major histocompatibility complex class I (MHC-I) immunopeptidome and increases neoantigen abundance to levels comparable to tumors with high mutational burden. This was associated with antigen-dependent T-cell-mediated tumor killing^23^. Despite this progress, it remains unclear which aberrant protein products are exposed at the cell surface after SMG1 inhibition. It remains unclear whether this remodeling is sufficient to elicit a measurable CD8⁺ T-cell effector response.

Identification of HLA-I-presented peptides arising from NMD-sensitive and noncanonical transcript isoforms requires expanding beyond canonical protein databases, which capture neither tumor-specific somatic mutations nor cell-specific, condition-dependent transcript diversity such as alternatively spliced isoforms and novel open reading frames. This can be achieved by integrating long-read RNA sequencing and ribosome profiling, which enables the generation of cell-specific protein sequence databases to identify peptides from these sources^31–35^. Moreover, pharmacological NMD inhibition stabilizes the potential neoantigen-encoding NMD-sensitive transcripts, making them available for translation and HLA-I presentation. Here, we applied this experimental approach in H838 NSCLC cells treated with the SMG1 inhibitor 11e (SMG1i).

HLA-I-bound peptides were identified by nanoflow liquid chromatography-tandem mass spectrometry (nLC-MS/MS) and further evaluated for compatibility with HLA-I binding using an extended metascore integrating NetMHCpan-derived HLA-I-binding predictions^36^ with NetChop-based proteasomal cleavage scores^37^, thereby incorporating two sequential steps of the HLA-I antigen-presentation pathway. Surface HLA abundance was assessed to distinguish changes in peptide composition from global alterations in HLA-I expression, while allogeneic CD8⁺ T-cell co-culture experiments evaluated the overall immunogenicity of SMG1i-treated cells rather than responses to individual selected peptides. Our findings suggest that pharmacological NMD inhibition reshapes the tumor immunopeptidome, revealing candidate antigenic peptides.

## Results

### SMG1 inhibition activates an NMD-perturbation signature in H838 NSCLC cells

To establish an experimental framework for subsequent immunopeptidomic analysis, H838 cells were treated with the SMG1i^29^ at different concentrations and exposure times to identify conditions suitable for subsequent immunopeptidomic analysis. Based on cell viability assays normalized to vehicle control, treatment with 0.2 µM and 0.3 µM SMG1i for 24 h retained approximately 82% and 55% metabolic activity, respectively, whereas 0.6 µM SMG1i reduced metabolic activity to approximately 41% (Supplementary Figure 1). Cells treated with 0.3 µM SMG1i for 8 h, a condition to assess early effects of SMG1 inhibition, retained 81% metabolic activity.

NMD activity was assessed by measuring the relative abundance of established NMD-sensitive and NMD-insensitive transcripts in SMG1i-treated and untreated cells. NMD-sensitive transcripts (*hnRNPL-NMD, RPSP,* and *TRA2B-NMD*) accumulated following SMG1 inhibition, with generally stronger increases (up to 60-fold) at higher inhibitor concentrations, whereas the NMD-insensitive control transcript of *TRA2B* remained unchanged (Figure 1.A). Selected UPF1 phosphopeptides included SQIDVALSQDSTYQGER with phosphorylation-site assignments at S1107, and AYQHGGVTGLSQY with an assignment at S1127.The UPF1-associated phosphopeptide signal in the phosphoproteomic analysis was lower on average following SMG1i treatment than in time-matched DMSO controls (Figure 1B). UPF1 protein was detected in all whole-cell proteome samples. Its abundance was higher on average following SMG1i treatment than in time-matched DMSO controls (Figure 1.C), opposite to its phosphopeptide intensities. Together, the transcript-level changes and label-free phosphoproteomic data confirmed a perturbation of the SMG1-UPF1-NMD axis.

**Figure 1.**
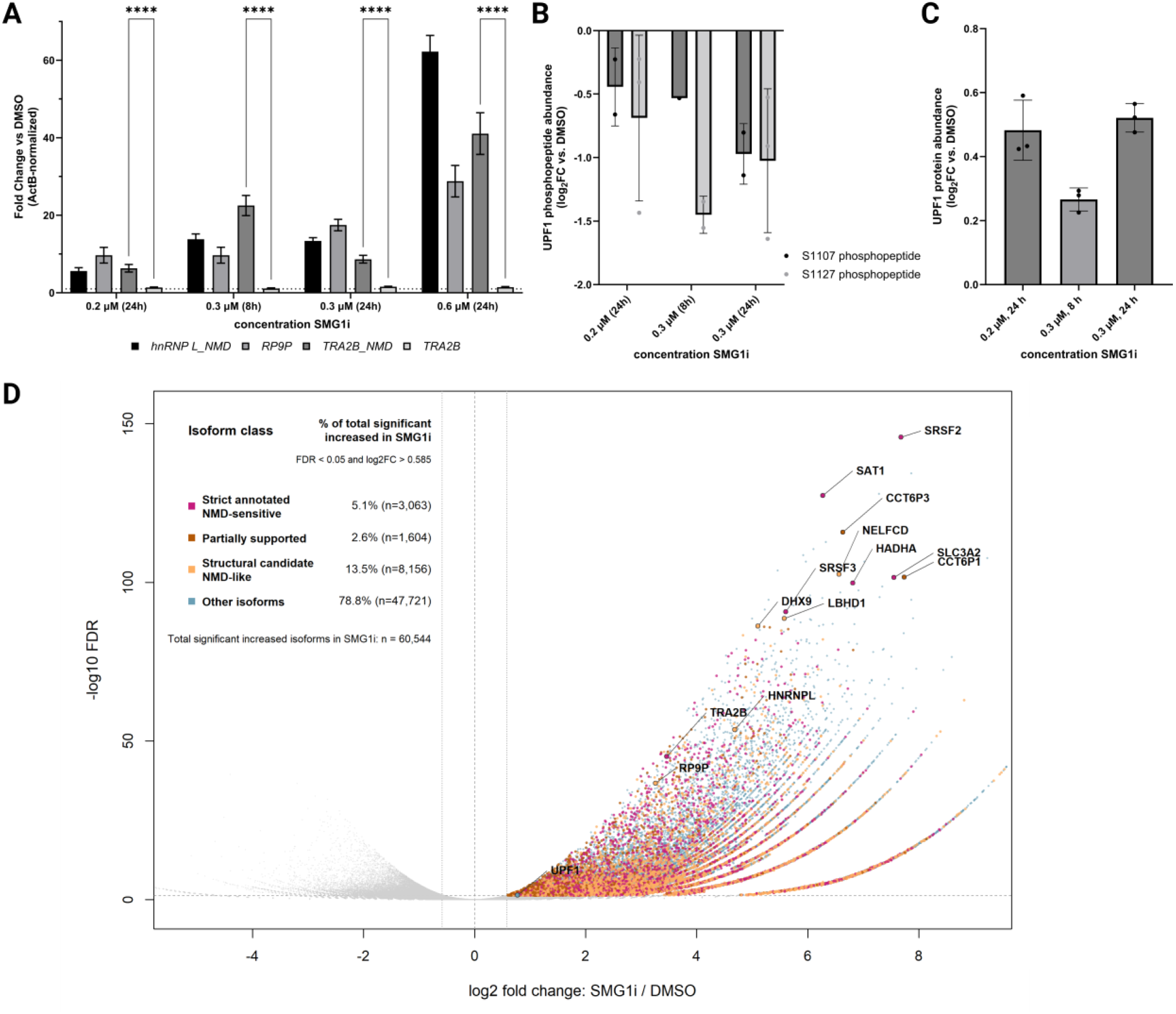
SMG1 inhibition activates an NMD-perturbation signature in H838 NSCLC cells. **(A**) H838 cells were treated with 0.2 µM SMG1i for 24 h, 0.3 µM SMG1i for 8 or 24 h, or 0.6 µM SMG1i for 24 h. Expression of the NMD-sensitive transcripts hnRNPL-NMD, RP9P, and TRA2B-NMD and the non-NMD TRA2B transcript was measured by RT-qPCR and normalized to ActB. Transcript abundance is expressed as fold change relative to the corresponding time-matched 0.1% DMSO control. Bars represent the mean ± SD of three technical replicates. Statistical significance was assessed by two-way repeated-measures ANOVA followed by Tukey’s multiple-comparisons test. ****adjusted p < 0.0001. **(B)** UPF1 phosphopeptide abundances measured by label-free nLC–MS/MS following treatment with 0.2 or 0.3 µM SMG1i for 24 h or 0.3 µM for 8 h. S1107- and S1127-containing phosphopeptides are shown in dark and light grey, respectively. Values are expressed as log₂FC relative to the mean log₂ quantity of available time-matched DMSO controls. Points represent individual treated replicates; bars show mean ± SD where calculable. In the order shown, n = 2, 1 and 2 for S1107 and 3, 2 and 3 for S1127. **(C)** UPF1 protein abundance measured by label-free nLC–MS/MS in the phospho-depleted fraction under the treatment conditions specified in (B). Each point represents the log₂ protein-group quantity of one treated replicate minus the mean log₂ quantity of time-matched DMSO controls. Bars show mean ± SD; n = 3 treated replicates per condition. **(D)** Differential isoform abundance following treatment with 0.6 µM SMG1i for 24 h compared with 0.1% DMSO was determined by PacBio Kinnex long-read RNA sequencing. The volcano plot shows log₂ fold change versus −log₁₀ FDR for all detected isoforms. Significantly upregulated isoforms (FDR < 0.05 and log₂FC > 0.585) are colored by NMD annotation category as strict annotated NMD-sensitive (magenta), partially supported NMD-like (brown), structural candidate NMD-like (yellow), and other isoforms (cyan). Selected NMD-related marker genes and highly significant isoforms are labeled. n = 3 technical replicates per condition.

Long-read RNA sequencing was used to assess transcript-isoform changes. Differential isoform analysis identified 60,544 significantly upregulated and 20,024 significantly downregulated transcript isoforms following SMG1 inhibition (false discovery rate (FDR) < 0.05 and |log₂FC| > 0.585; Figure 1.D). Among the upregulated isoforms, one UPF1 transcript isoform was also significantly increased following SMG1 inhibition (log₂FC = 0.77, FDR = 0.0417). Of the 60,544 upregulated isoforms, 3,063 carried a GENCODE NMD annotation and/or transcript-level NMDtxDB evidence combining a premature termination codon with significant accumulation following NMD-factor perturbation^38,39^. Of these 3,063 NMD-annotated upregulated isoforms, log₂FC ranged from 0.66 to 8.62, with a median of 3.64. NMD annotations were detected in 5.06% of SMG1i-upregulated isoforms compared with 1.55% of the remaining 684,372 analyzed isoforms that did not meet the upregulation criteria, corresponding to an odds ratio of 3.38 (95% confidence interval, 3.24-3.52).

### Immunopeptidomics coupled to proteogenomics identifies a high-confidence HLA-I-associated peptide repertoire

To identify HLA-I-bound peptides beyond the canonical proteome, a H838-specific proteogenomic sequence database was generated from PacBio Kinnex long-read RNA-seq and Illumina ribo-seq data generated from H838 cells with and without SMG1i treatment. The resulting proteogenomic sequence database was concatenated with the human UniProtKB reference proteome (UP000005640, release 2025_12)^40^. Known MS contaminant sequences were also included to prevent false matches of such peptides. Entries with an appended UniProtKB entry were classified as cell-specific and canonical, whereas entries without were retained as cell-specific only.

Using the four SMG1i treatment conditions defined above, with matched vehicle control samples, were analyzed in triplicate, except for the 0.6 µM condition, which was analyzed in six replicates. In total, we identified 12,352 peptide forms. Application of the calculated extended metascore cutoff, being 0.0600 for a 1% FDR, retained 2,728 peptides (Figure 2.A).

**Figure 2.**
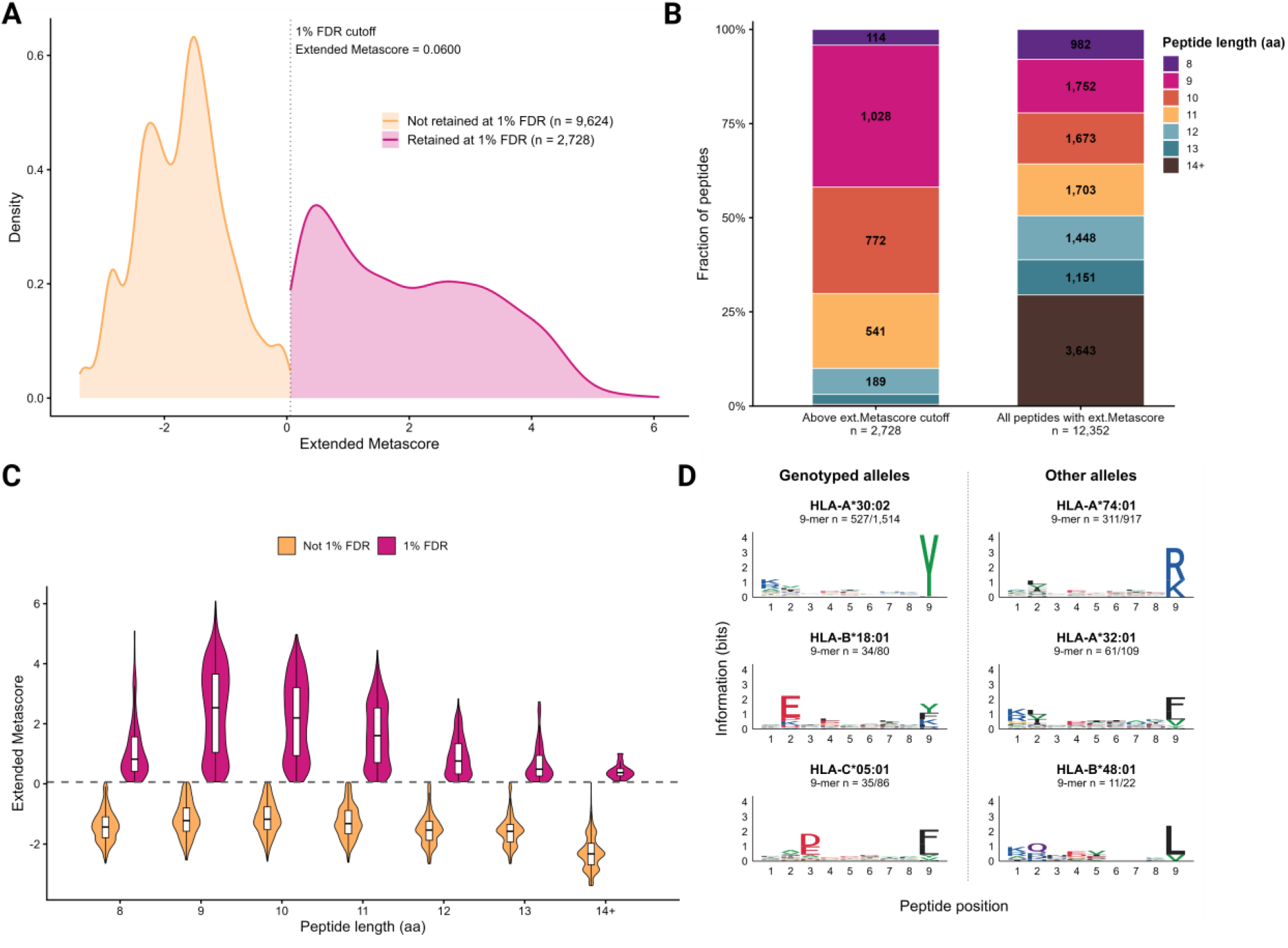
Extended-metascore filtering and characteristics of retained HLA class I peptide identifications. **(A)** Distribution of extended metascores for all peptide identifications. Yellow and magenta indicate identifications below and at or above the score threshold corresponding to 1% peptide-level FDR, respectively. The vertical dotted line marks this threshold. Smoothed density curves are clipped at the threshold. **(B)** Peptide-length distribution of potentially true versus all peptides based on the extended metascores 1% FDR cutoff. **(C)** Extended-metascore distributions of peptide identifications stratified by peptide length. Peptides retained at 1% FDR (magenta) are separated from those not retained (yellow). Embedded boxplots indicate the median and interquartile range. The dashed horizontal line marks the extended metascore of cutoff value of 0.0600. **(D)** HLA class I peptide motifs of 1% FDR 9-mers. Sequence logos show the motifs of unique 9-mer peptide sequences of the potentially true binders, grouped by their best-predicted HLA assignment. The number above each panel indicates the number of unique 9-mer sequences contributing to the motif.

The 1% FDR-filtered population was strongly enriched for peptide lengths typical for HLA-I ligands, with 9-, 10-, and 11-mer peptides accounting for 38%, 28%, and 20% of the entries, respectively, whereas longer peptides were markedly depleted (Figure 2.B). High-resolution HLA genotyping identified HLA-A*30:02, HLA-B*18:01, and HLA-C*05:01 in H838 cells. Predicted peptide-to-HLA assignments were generated using NetMHCpan^36^ with a six-allele prediction panel comprising the genotyped H838 alleles together with HLA-A*32:01, HLA-A*74:01, and HLA-B*48:01. The three additional alleles included in the prediction panel were not detected by HLA genotyping and were used only for computational peptide assignment. HLA-B*48:01 was included because it was reported in the Cellosaurus NCI-H838^41,42^ entry based on previously published HLA-typing data^41,42^. The binding motif of HLA-A*74:01 has a similarity to a reversed motif of HLA-A*30:02, and the HLA-B*48:01 motif can be regarded as a competitor for HLA-B*18:01.

The extended metascore distributions of the predominant 9-11-mer population showed a clear bimodality (Figure 2.C). The higher-scoring population containing stronger predicted binders was predominantly assigned to HLA-A*30:02 (56% of all peptides) and had greater peptide-spectrum match (PSM) support than the lower-scoring population, with a median number of PSMs per peptide length class between 10-15 compared with 2-3 in the lower-scoring peptide population. Furthermore, the lower-scoring population contained a large proportion of HLA-A*74:01 assignments.

Motif analysis of the 1% FDR-filtered 9-mer peptides revealed distinct position-specific residue preferences consistent with predicted HLA-I binding motifs (Supplementary Figure 2). HLA-A*30:02-assigned peptides showed a prominent C-terminal tyrosine preference, whereas peptides assigned to HLA-A*74:01 displayed a distinct C-terminal arginine/lysine pattern, as expected from their respective predicted binding motifs (Figure 2.D).

Application of the 1% FDR threshold to the extended metascore defined a high-confidence HLA-I peptide set. The expected characteristics of the retained peptides, including their predominant assignment to the genotyped HLA-A*30:02 allele, supported both the biological plausibility of the selected peptide repertoire and the use of the extended metascore framework for subsequent treatment-associated analyses.

### SMG1 inhibition remodels the HLA-I immunopeptidome without increasing HLA-I surface expression

When grouped by extended metascore, vehicle controls contributed more PSMs in the lower-scoring range, whereas SMG1i-treated samples contributed more PSMs within the high-confidence range (Figure 3.A). In contrast, the numbers of unique peptide sequences were comparable between conditions within the high-confidence range (Figure 3.B). Together, these findings indicate increased abundance for a restricted peptide subset following SMG1 inhibition rather than an expansion of the HLA-I-presented peptide repertoire.

**Figure 3.**
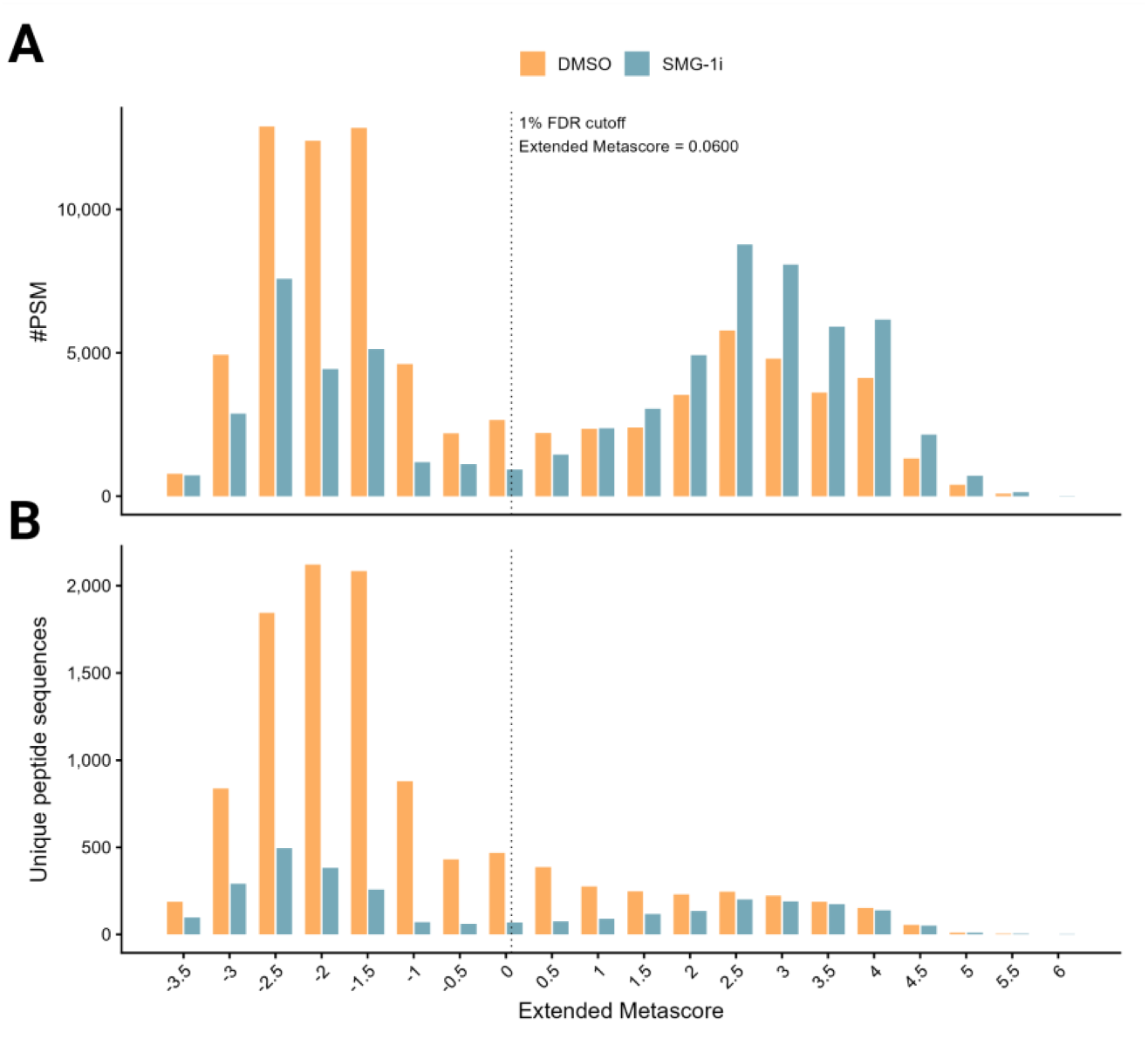
Treatment-associated peptide evidence and prioritization following SMG1 inhibition in H838 cells. **(A, B)** Peptide identifications from DMSO- and SMG1i-treated H838 samples were grouped into 0.5-unit extended-metascore bins. **(A)** Aggregated peptide-spectrum match (PSM) counts within each bin. **(B)** Number of unique peptide sequences within each bin. The vertical dotted line indicates the extended-metascore cutoff corresponding to 1% peptide-level FDR.

Of the 2,728 peptides retained as high-confidence predicted HLA-I binders, 1,310 (48.0%) had insufficient PSM support for assignment to treatment-response(see Methods), leaving 1,418 peptides. Among these, 696 (49%) were unchanged between untreated and SMG1i-treated cells, 48 (3%) were detected only in SMG1i-treated cells, and 273 (19%) were enriched upon SMG1i incubation, whereas 339 (24%) and 62 (4%) were only detected and enriched in untreated cells, respectively. The data support therefore a remodeling effect of the HLA-I immunopeptidome upon SMG1i treatment. Therefore, we focused (see also next section) on the 48 peptides that were exclusively detected in the SMG1i-treated cells.

Of these, 37 (77%) matched both cell-specific and canonical protein sequences, 5 (10%) matched cell-specific sequences only, and 4 (8%) matched canonical sequences. Two peptides (4%) were identified as known MS contaminants and were excluded from further analysis.

To determine whether the appearance of SMG1i-induced peptides was accompanied by increased surface HLA-I abundance, surface expression of HLA-I was assessed with the conformation-dependent anti-HLA-I W6/32 and an isotype control antibody by flow cytometry. Nearly all gated H838 cells were W6/32-positive under all treatment conditions. Surface HLA-I abundance was therefore quantified as the median AF647-A fluorescence of gated singlet cells. W6/32 and isotype-control fluorescence values were compared on the same scale (Figure 4.A). SMG1 inhibition did show a decrease in W6/32 fluorescence with both SMG1i concentrations and antibody dilutions used. A relatively smaller decrease was also detected in the corresponding isotype controls, whose absolute fluorescence was similar at both antibody dilutions. This nonspecific shift may reflect broader treatment-associated effects consistent with the previously observed reduction in metabolic activity. However, cell loss alone cannot explain the result, because fluorescence was quantified as the median signal of individual gated singlet cells rather than as total sample fluorescence. The W6/32 measurements therefore provided no evidence of increased HLA-I surface abundance, while the contribution of nonspecific treatment effects prevented the observed decrease from being interpreted as definitive HLA-I downregulation. Representative immunofluorescence images were consistent with the absence of increased surface HLA-I staining following SMG1 inhibition (Figure 4.B).

**Figure 4.**
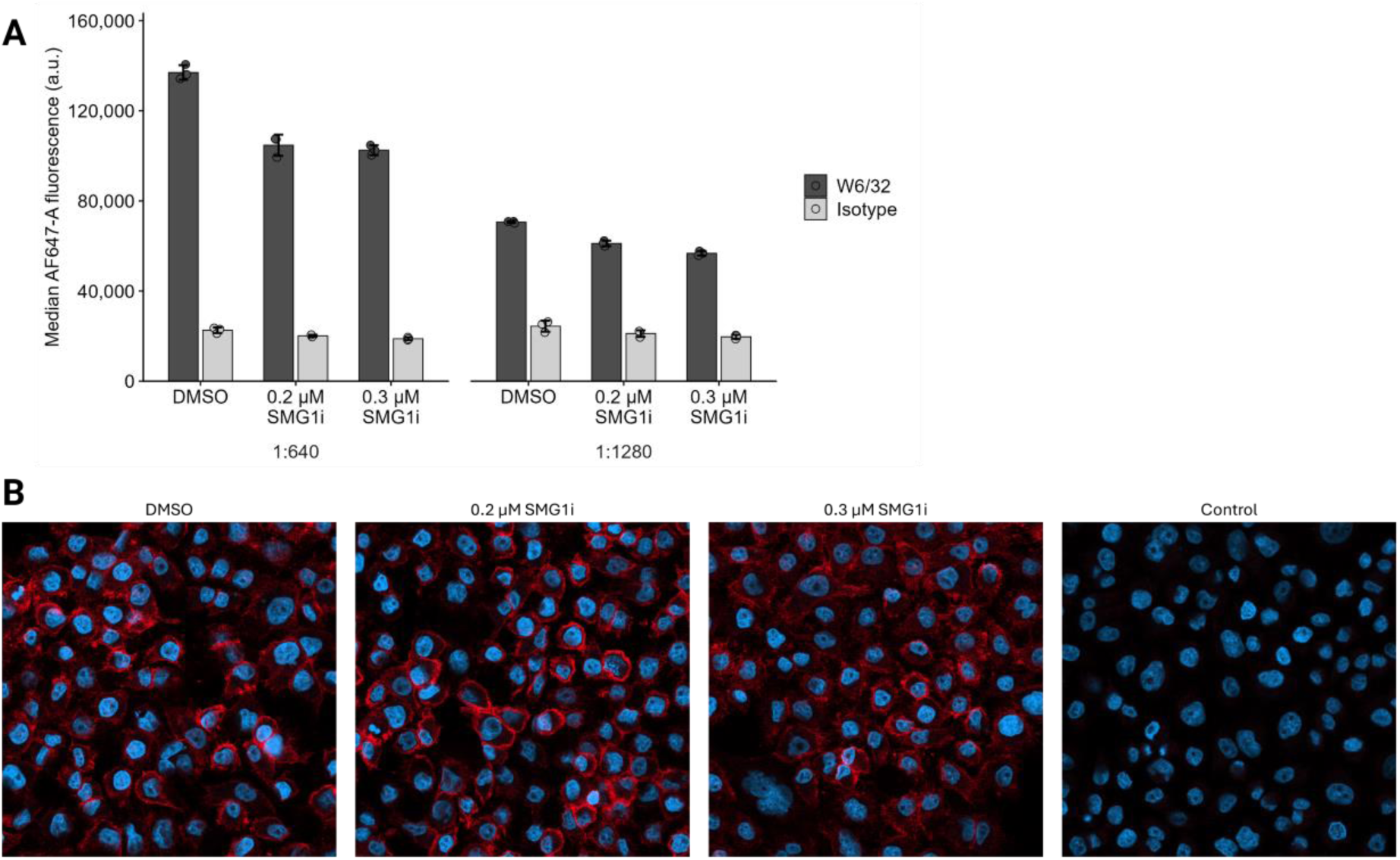
SMG1 inhibition does not increase surface HLA on H838 cells. **(A)** Flow-cytometric quantification of surface HLA class I on H838 cells treated with DMSO, 0.2 µM SMG1i, or 0.3 µM SMG1i for 24 h and stained with W6/32 antibody at 1:640 and 1:1280 dilutions. Raw median AF647-A fluorescence of gated singlet cells is shown for W6/32 and isotype controls on the same scale. Individual points represent technical replicate measurements and bars indicate the mean ± SD. No statistical testing was performed. a.u. stands for arbitrary units. **(B)** Representative immunofluorescence images of H838 cells after 24 h treatment with DMSO, 0.2 µM SMG1i, or 0.3 µM SMG1i. Cells were formalin-fixed without permeabilization and stained with W6/32 conjugated to Alexa Fluor 647 to detect HLA class I (red). Nuclei were counterstained with DAPI (blue). The isotype-control image shows background staining with the corresponding Alexa Fluor 647-conjugated mouse IgG2a antibody.

### Functional annotation of newly presented peptides

The associated proteins of the 32 SMG1i treatment-specific immunopeptides (Supplementary Table 1) represented diverse cellular processes, including translation, RNA processing and protein quality control, metabolism and cofactor homeostasis, organelle homeostasis and intracellular trafficking, and genome maintenance, respectively.

Peptides assigned to sequences of NAA16, MRRF, and WASH-family genes were detected across all SMG1i concentrations. NAA16 and MRRF are involved in co-translational processing and mitochondrial ribosome recycling, linking this subset to translation-related protein quality control. In contrast, WASH-family proteins regulate endosomal actin nucleation and cargo sorting, and the corresponding peptide mapped to several related sequences, including pseudogene-annotated loci, preventing assignment to a single source gene and warranting cautious interpretation of its source attribution. This ambiguous source attribution nevertheless made the peptide of interest because noncanonical translation products can preferentially contribute to the HLA-I immunopeptidome.

Beyond this recurrent subset, other peptides were assigned to ACSS1, PGM1, and PTS, involved in metabolic and cofactor homeostasis, FASN (lipid synthesis and membrane remodeling), RPL13A (translational control), TIMM13 (mitochondrial protein import), and ATR (replication-stress signaling and genome maintenance)_40_.

Two peptides assigned to DHCR24 and CLSTN1, were found only in the samples treated with 0.2 µM and 0.3 µM SMG1i and were associated with cholesterol and oxidative-stress homeostasis and vesicular trafficking and membrane dynamics, respectively.

Peptides assigned to EPG5, LMBRD1, GUSB, MMS19, RBM39, PUM1, GIPR, HSP70-family proteins, DARS1, and IFT52 were restricted to the 0.6 µM SMG1i condition. EPG5, LMBRD1, and GUSB represented autophagosome-lysosome fusion, lysosomal transport, and lysosomal degradation, respectively, while RBM39 and PUM1 were associated with RNA processing and post-transcriptional regulation. HSP70-family proteins and DARS1 linked this subset to protein folding and translation-related homeostasis, whereas MMS19 was associated with genome maintenance, GIPR with metabolic signaling, and IFT52 with ciliary transport. The restriction of this peptide subset to the 0.6 µM condition coincided with a reduction in metabolic activity to approximately 41% (see “SMG1 inhibition activates an NMD-perturbation signature in H838 NSCLC cells”), indicating that 0.6 µM SMG1i treatment triggered pronounced cellular stress responses.

The immunogenic potential of these peptides was predicted using PRIME 2.1^43^. Additional prediction approaches provided complementary information on immunogenic potential and peptide-HLA-I complex stability^44–46^. Lower PRIME percentile ranks indicate stronger predicted immunogenic potential relative to the model’s reference peptide population^43^. Five recurrent HLA-A*30:02-assigned candidates were examined in more detail based on their spectral support. Among these, RVPGIRSHLY received the strongest combined computational support, with a PRIME percentile rank of 0.034% and a predicted peptide-HLA-I complex half-life of 2.95 h. This peptide was assigned to an MRRF-associated sequence, detected with 20 PSMs across 11 of 15 SMG1i-treated samples, and matched both cell-specific and canonical protein sequences. RVKVFQSSANY also showed a favorable PRIME prediction, with a percentile rank of 0.318%, and the highest predicted peptide-HLA-I complex stability among the five candidates, with a half-life of 4.41 h. AQHFDKLGQY showed a favorable PRIME percentile rank of 0.232% but lower predicted complex stability, with a half-life of 0.47 h. RMQHSLAGQTY and RISFSASHRLY received more moderate computational support, with PRIME percentile ranks of 0.786% and 0.816% and predicted complex half-lives of 2.74 and 1.38 h, respectively. TIDDGIFEV and VGDIPGVRF were the only final candidates assigned to HLA-C*05:01 rather than HLA-A*30:02 and received favorable allele-specific PRIME percentile ranks of 0.005% and 0.010%, with peptide-HLA-I complex half-lives of 0.19 h for TIDDGIFEV and 0.09 h for VGDIPGVRF. Together, the immunogenicity and peptide-HLA-I stability predictions provided additional criteria for comparing the candidates but did not establish their immunogenicity experimentally.

### SMG1i treatment is associated with enhanced CD8⁺ T-cell responses

To assess whether SMG1 inhibition activated cell-intrinsic innate immune signaling that could contribute to T-cell responses, the expression of the representative interferon-stimulated genes (ISGs) *IFI44L, ISG15*, and *MX1*^47–52^ was measured in SMG1i-treated and vehicle-treated H838 cells. None of these ISGs were upregulated following SMG1 inhibition. *IFI44L* and *ISG15* expression remained unchanged, whereas *MX1* expression was modestly downregulated, providing no evidence for induction of a canonical type I interferon transcriptional response under the tested conditions (Figure 5.A). These findings indicated that SMG1 inhibition did not trigger detectable type I interferon responses in H838 cells, which could potentially contribute to T-cell activation.

**Figure 5.**
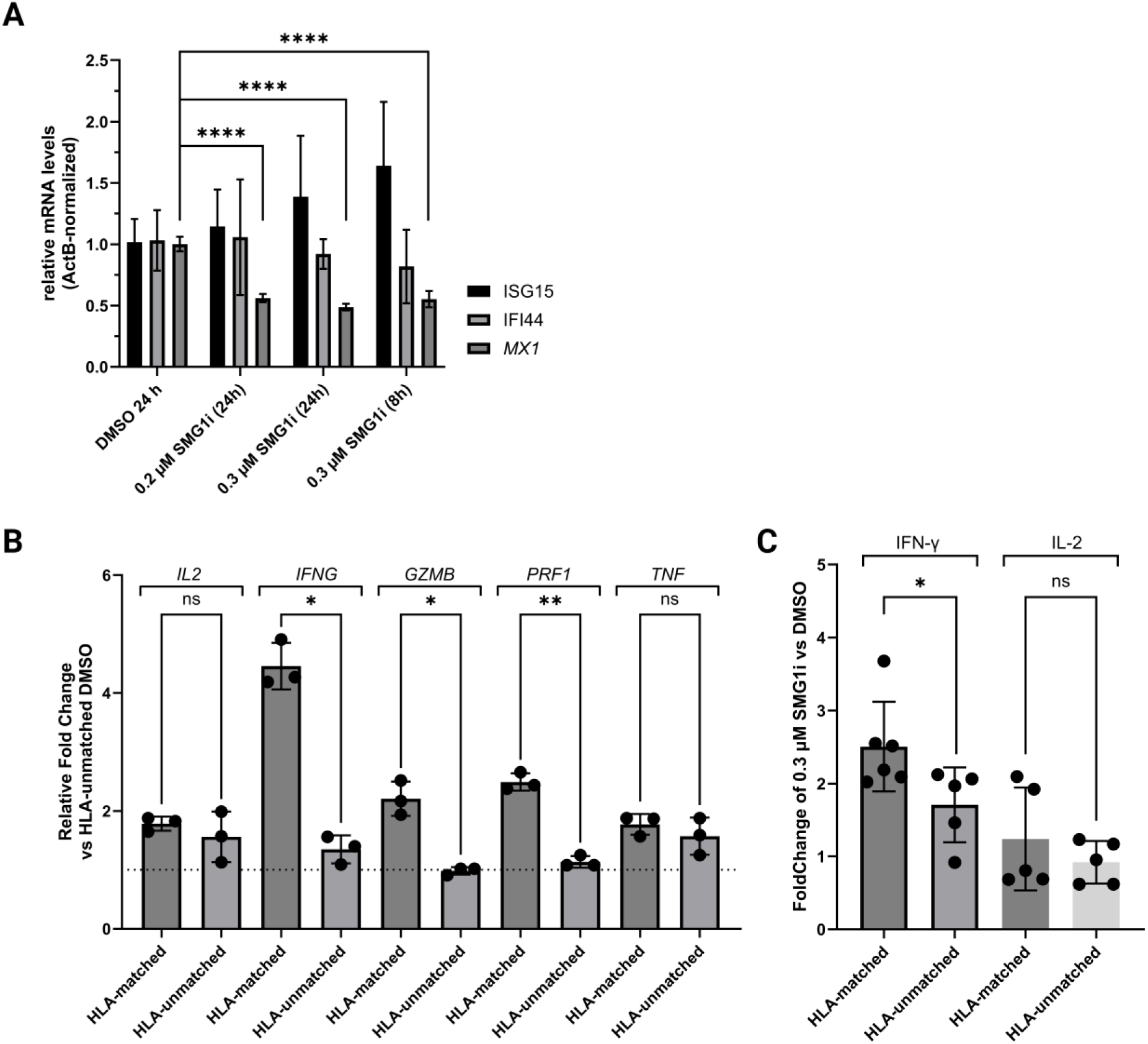
SMG1 inhibition does not induce an interferon response in H838 cells but is associated with enhanced CD8⁺ T-cell responses in HLA-matched co-cultures. **(A)** Expression of ISG15, IFI44, and MX1 was measured by RT-qPCR following treatment of H838 cells with DMSO, 0.2 µM SMG1i for 24 h, or 0.3 µM SMG1i for 8 or 24 h. Comparative threshold-cycle (Ct) values were normalized to ACTB and expressed relative to the DMSO control. Bars show the mean ± SD of three experiments. Statistical significance was assessed separately for each transcript by one-way repeated-measures ANOVA followed by Dunnett’s multiple-comparisons test against DMSO. **(B)** H838 cells pretreated with 0.3 µM SMG1i or DMSO for 24 h were co-cultured with isolated and activated CD8⁺ T cells from one HLA-matched and one HLA-unmatched donor. Expression of IL2, IFNG, GZMB, PRF1, and TNF was measured by RT-qPCR. Bars show the mean ± SD of three experiments and dots show individual measurements. **(C)** H838 cells pretreated with 0.3 µM SMG1i or DMSO for 24 h were co-cultured with isolated and activated CD8⁺ T cells from fresh or cryopreserved PBMCs of HLA-matched (n = 2 donors) or HLA-unmatched (n = 1 donor) donors. Intracellular IFN-γ and IL-2 were measured by flow cytometry and expressed as fold change relative to the corresponding donor-matched DMSO control. Bars show the mean ± SD and dots show individual measurements. HLA-matched and HLA-unmatched groups were compared separately for each cytokine using two-sided unpaired Welch’s t-tests. *p < 0.01, **p < 0.001, ****p < 0.0001; ns, not significant.

Because autologous immune cells were not available for the established H838 cell line, an allogeneic co-culture model was used to assess whether SMG1i-treated H838 cells enhanced CD8⁺ T-cell responsiveness. Two donors could be recruited, each sharing one allele at HLA-A, HLA-B, and HLA-C with H838, corresponding to a 3/6 allelic match, enabling potential recognition of H838-presented peptides. A donor without any shared alleles served as the unmatched control.

H838 cells were pre-treated with 0.3 µM SMG1i or vehicle and exposed to primary CD8⁺ T-cells from non-cryopreserved PBMCs from of the partially HLA-matched donors and the unmatched donor. Low but detectable IFN-γ responses were observed by flow cytometry, with approximately 2-3% of CD8⁺ T-cells from the partially HLA-matched donor compared with approximately 1.3-1.5% from the unmatched donor. IL-2-positive cells remained below 1% across all co-culture conditions. To assess cytokine and cytotoxic effector responses, RT-qPCR analysis was performed for the cytokine effector genes *IFNG* and *TNF*, as well as components of the cytotoxic machinery, *GZMB* and *PRF1,* and *IL2* as a cytokine associated with T-cell proliferation. Consistent with the flow-cytometry results, *IFNG*, *GZMB*, and *PRF1* showed significant induction in the partially HLA-matched donor compared with the unmatched one, while no significant increase in *IL2* and *TNF* mRNA production was detected (Figure 5.B).

Additionally, data obtained using cryopreserved PBMCs from all three donors (n = 3; two partially HLA-matched, one unmatched) were combined with the non-cryopreserved PBMCs experiments (Figure 5.C). Intracellular IFN-γ levels were significantly higher in both partially HLA-matched donors than in the unmatched donor. No effect on IL-2 was confirmed. Given the limited donor number (n = 3; two partially matched, one unmatched), this was interpreted as an exploratory trend indicating HLA-dependent activation of CD8^+^ T-cells.

## Discussion

Here, we investigated whether inhibition of SMG1 with a small-molecule inhibitor and the resulting attenuation of NMD remodels the HLA immunopeptidome of H838 NSCLC cells and is associated with enhanced CD8⁺ T-cell responses. Across the different molecular layers examined, SMG1 inhibition reduced the degradation and increased the abundance of NMD-sensitive transcript isoforms, altered the transcript and translation landscape, was associated with a lower UPF1-associated protein-group signal in the phosphoproteomic analysis, and remodeled HLA-I peptide presentation without an apparent change in HLA presentation. The latter findings were supported by a stable surface HLA abundance measured by flow cytometry and immunofluorescence staining, while the number of unique peptide sequences was comparable between SMG1i- and vehicle-treated samples, but with concordant treatment-associated changes in the peptide repertoire. Finally, co-culture experiments showed enhanced CD8⁺ T-cell responses in the partially HLA-matched donor setting, including increased IFN-γ and induction of effector-associated transcripts.

Genotyping showed that H838 is homozygous at the HLA-A, HLA-B, and HLA-C loci. Concordant results were obtained using two sequencing-based HLA-typing approaches, Illumina short-read and Oxford Nanopore sequencing. The three additional alleles included in the NetMHCpan prediction panel, HLA-A*32:01, HLA-A*74:01, and HLA-B*48:01, were not detected by either genotyping approach. They were included because of similarities in predicted binding motifs, previously reported H838 HLA typing compiled in Cellosaurus^41,42^, and as additional motif proxies for calculation of the false discovery rate cutoff. The additional alleles were therefore used only for computational peptide assignment and should not be interpreted as evidence of their expression in H838 cells. The observed homozygosity may reflect either germline HLA homozygosity or somatic loss of heterozygosity (LOH). Germline HLA-I homozygosity can itself restrict the repertoire of potentially immunogenic peptides available for presentation^53^, whereas somatic HLA LOH represents a recurrent immune-evasion mechanism in NSCLC that can further reduce the breadth of tumor antigen presentation^54^.

SMG1-dependent phosphorylation of UPF1 is a central step in mammalian NMD activation and is required for efficient degradation of NMD-targeted transcripts^55,26^. Compound 11e was originally described as a potent pyrimidine-derived SMG1 inhibitor with an improved kinase-selectivity profile^29^. Consistent with NMD inhibition, treatment of H838 cells with compound SMG1i 11e was associated with a decreased UPF1 phosphopeptide signal and an increased abundance of well-characterized NMD-sensitive transcripts, including the NMD-sensitive *HNRNPL* and *TRA2B* isoforms and *RPSP*^56^. Although the UPF1 phosphoproteomic data overall support SMG1 target engagement, limited site coverage precluded robust site-specific conclusions. In addition, because the phosphoproteomic and proteomic measurements were acquired separately, the UPF1 phosphopeptide signal could not be directly normalized to the total UPF1 protein signal. Total UPF1 protein signal showed a trend in the opposite direction to the UPF1-associated protein-group signal in the phosphoproteomic analysis, and one UPF1 transcript isoform was significantly increased following SMG1 inhibition. This pattern is consistent with reduced UPF1 phosphorylation occurring alongside a trend toward increased UPF1 abundance, in agreement with the established autoregulation of the NMD pathway following NMD inhibition^57^. Of the detected phosphorylation sites, S1127 is located within an S/TQ motif preferentially recognized by PIKK-family kinases, including SMG1. However, the presence of this motif does not by itself demonstrate that SMG1 directly phosphorylates these residues. Nevertheless, the combined phosphoproteomic, proteomic, and transcriptomic observations consistently support the perturbation of the SMG1-UPF1-NMD axis by SMG1i 11e and provide a mechanistic basis for interpreting the downstream molecular and immunopeptidomic changes.

Because NMD inhibition prevents the efficient degradation of NMD-sensitive transcript isoforms that may subsequently be translated, it can expand the sequence space available for HLA-I peptide generation. A key feature of the immunopeptidomic analysis was therefore the use of a cell-specific database generated from long-read RNA-seq and ribo-seq data obtained from treated and untreated H838 cells^31,58,59^. Annotation of the H838-derived sequences against UniProtKB enabled the distinction between sequences with a corresponding canonical protein match and sequences supported exclusively by the cell-specific transcriptomic and translatomic data.

Within this large sequence search space, the extended metascore provided an additional mathematical framework to prioritize candidate HLA-I ligands beyond statistical identification confidence by fragment spectra interpretation software alone. By integrating NetMHCpan-derived HLA-binding predictions with NetChop-based proteasomal cleavage predictions, the score considered two complementary determinants that constrain HLA-I antigen presentation^36,37^. The combined use of predicted HLA-I binding with upstream antigen-processing information is consistent with established epitope-prioritization approaches and can improve candidate selection compared with HLA-I-binding prediction alone^58,60^. In the present analysis, these predictions were complemented by experimental evidence including semi-quantitative spectrum counts and recurrent detection across many SMG1i-treated samples, thereby favoring reproducible candidates compatible with the H838 HLA-I genotype. The predicted immunogenic potential of the prioritized candidates was subsequently assessed using PRIME 2.1, complemented by peptide-HLA-I complex-stability predictions using NetMHCstabpan^43,46^. These analyses refined their relative prioritization but did not experimentally establish peptide-specific T-cell recognition.

Among the prioritized peptides, the WASH-family candidate was particularly interesting because its sequence also mapped to pseudogene-annotated loci. Pseudogenes and other noncanonical transcripts can contribute translated products to the HLA-I immunopeptidome^61^, while SMG1 inhibition has been shown to expose non-mutational, noncanonical antigenic sequences normally constrained by NMD^23^. However, its ambiguous mapping prevents assignment to a defined transcript of origin and therefore precludes its classification as a neoantigen without further validation.

1Furthermore, five recurrent HLA-A*30:02-assigned candidates were examined in greater detail based on their experimental support. Of these, RVPGIRSHLY received the strongest combined computational support. Only two prioritized peptides were assigned to HLA-C*05:01. Their limited representation may partly reflect the lower surface expression and more restricted peptide-binding repertoire reported for HLA-C relative to HLA-A/B, consistent with the short predicted peptide-HLA-I complex half-lives observed for these candidates^62–64^.

Two further considerations constrain this prioritized set. First, candidates restricted to 0.6 µM SMG1i require more cautious interpretation because this condition coincided with markedly reduced metabolic activity and may therefore reflect broader cellular stress in addition to consequences of NMD inhibition. Second, peptides designated “SMG1i-only” were detected in treated samples but not in vehicle controls. This does not establish their absence from untreated cells, where their abundance may have been below the detection limit. Importantly, these peptide-level changes occurred without a corresponding increase in total surface HLA-I. Because W6/32 detects conformational HLA-I complexes collectively rather than individual peptide-HLA-I species, the absence of increased global HLA-I abundance indicates a substantial remodeling of the presented peptide repertoire^65,66^.

The response in partially HLA-matched primary CD8⁺ T-cell co-cultures suggests that HLA compatibility contributed to the recognition of SMG1i-treated H838 cells, although peptide-specific recognition was not established. Increased IFN-γ production together with concordant induction of *IFNG, GZMB*, and *PRF1* is consistent with enhanced CD8⁺ T-cell effector activity following SMG1 inhibition^67^. The absence of a corresponding IL-2 response does not necessarily indicate failed CD8⁺ T-cell activation, as IL-2 production depends more strongly on additional co-stimulatory signals and CD4⁺ T-cell help than cytotoxic effector functions such as IFN-γ production^68,69^. A separate H838 monoculture experiment provided no evidence for generalized cell-intrinsic type I interferon activation following SMG1 inhibition, as *IFI44L* and *ISG15* were not induced, and *MX1* was modestly reduced. This makes such an innate response an unlikely primary explanation for the enhanced T-cell response observed in co-culture.

The allogeneic nature of the co-culture nevertheless limits mechanistic interpretation of this response. HLA compatibility between H838 and the partially matched donors is consistent with an HLA-dependent contribution, but residual alloreactivity and other treatment-induced changes in the tumor cells cannot be excluded. Consequently, the enhanced response cannot yet be attributed to individual SMG1i-associated peptide-HLA-I complexes. Direct tumor-cell killing was also not assessed. Peptide-specific validation will therefore be required to determine whether the prioritized HLA ligands themselves contribute to recognition of SMG1i-treated H838 cells.

Our findings are consistent with emerging evidence that pharmacological SMG1 inhibition can reshape tumor antigen presentation and enhance immune recognition. Cook and colleagues showed increased HLA-I presentation of peptides derived from NMD-regulated proteins following treatment with the *in vivo*-compatible SMG1 inhibitor KVS0001^25^. Vendramin and colleagues further demonstrated broader immunopeptidome remodeling, increased neoantigen presentation, and antigen-dependent T-cell-mediated tumor killing following SMG1 inhibition^23^. The translational relevance of SMG1 inhibition is underscored by the first-in-human evaluation of DS5361, alone and in combination with pembrolizumab, in patients with TMB-high and/or MSI-high advanced solid tumors^70^. While these populations are enriched for mutation-derived neoantigens, the present data suggest that NMD inhibition can also expose antigens of non-mutational origin. The treatment-associated candidates identified here provide a complementary peptide space in which this broader antigenic potential can be investigated.

Future work should focus on confirming the identity and functional relevance of the prioritized SMG1i-associated peptides. The five HLA-A*30:02 strong-binder candidates highlighted here should first be validated against synthetic peptide standards, followed by peptide-specific CD8⁺ T-cell priming and testing of peptide-reactive T-cells against SMG1i-treated and vehicle-treated H838 cells. This will determine whether peptide-specific CD8⁺ T-cells recognize SMG1i-treated H838 cells.

Beyond H838, extending this approach to NSCLC models with distinct HLA backgrounds and ultimately to patient-derived tumors paired with autologous immune cells will be important for assessing the generalizability and translational relevance of SMG1-dependent immunopeptidome remodeling. Together, the present study shows that pharmacological SMG1 inhibition can reshape the HLA-I peptide repertoire in an NSCLC model and identifies a set of treatment-associated peptide candidates for functional validation.

## Methods

### Cell Culture and SMG1 Inhibitor Treatment

NCI-H838 cells were cultured in Dulbecco’s modified Eagle medium (DMEM, Gibco, Cat. No. 41965-039) supplemented with 10% fetal bovine serum (FBS; Gibco, Cat. No. A5256801), 100 U/mL penicillin, and 100 µg/mL streptomycin at 37 °C in a humidified atmosphere containing 5% CO₂. HB-95 hybridoma cells used for W6/32 antibody production were maintained under the same conditions. NCI-H838 cells were treated with 0.1 µM to 0.6 µM SMG1 inhibitor 11e (SMG1i)^29^ or 0.1% dimethyl sulfoxide (DMSO) as vehicle control. Treatment duration depended on the experiment and is specified in the corresponding experimental sections. For immunopeptidomic analysis, cells were harvested in ice-cold Dulbecco’s phosphate-buffered saline (DPBS) and centrifuged at 500 × g for 5 min at 4 °C. Cell pellets corresponding to approximately 1 × 10⁷ cells were flash-frozen in liquid nitrogen and stored at −80 °C until enrichment of peptide-HLA-I complexes.

### HLA Genotyping

Genomic DNA was isolated from untreated NCI-H838 cells and donor-derived peripheral blood mononuclear cells (PBMCs) using the Maxwell RSC Cultured Cells DNA Kit (Promega, Cat. No. AS1620) according to the manufacturer’s instructions.

HLA genotyping was performed using the IVD-certified AllType FASTplex NGS 11 Loci Flex Kit (One Lambda, Thermo Fisher Scientific, Cat. No. ALL-FAST11LF) on an Illumina iSeq 100 system. The assay covered HLA-A, HLA-B, HLA-C, HLA-DR, HLA-DQ, and HLA-DP loci. Sequence data were analyzed using TypeStream Visual NGS Analysis Software version 3.1 (One Lambda, Thermo Fisher Scientific), with analysis engine version 3.1.0.58 and IPD-IMGT/HLA database version 3.60.0.0.

The HLA genotype of NCI-H838 cells was independently confirmed by long-read HLA sequencing using the CE-IVD NanoTYPE 96/11 Kit (Omixon Biocomputing Ltd., Cat. No. NT9611CE) on an Oxford Nanopore MinION platform. Reads were processed using MinKNOW version 22.5, and HLA alleles were assigned using NanoTYPER RUO version 2.1.0 against IMGT/HLA reference database version 3.60.0_10. Allele assignments from both sequencing approaches were reviewed by the “Zentrum für Labormedizin” (Insel, Bern) before reporting. HLA-I compatibility between H838 cells and PBMC donors was determined by comparing the assigned HLA-A, HLA-B, and HLA-C alleles.

### Cell Viability Assay

H838 cell viability was assessed using the 3-(4,5-dimethylthiazol-2-yl)-2,5-diphenyltetrazolium bromide (MTT, Sigma-Aldrich, Cat. No. 298-93-1) assay. Cells were treated with 0.1-10 µM SMG1i or 0.1% DMSO vehicle control for 1-24 h. Following treatment, cells were incubated with 0.5 mg/mL MTT for 3 h at 37 °C. Formazan crystals were dissolved in DMSO, and absorbance was measured at 570 nm using a Sunrise microplate reader (Tecan). Cell viability was expressed relative to the corresponding DMSO vehicle control.

### RNA Isolation, Reverse Transcription, and RT-qPCR

Total RNA was isolated using the Maxwell RSC simplyRNA Cells Kit (Promega, Cat. No. AS1390) according to the manufacturer’s instructions. Relative transcript abundance was calculated using the comparative threshold-cycle method ( 2^−ΔΔ*Ct*^)^71^. Primer sequences for all reverse-transcription quantitative PCR (RT-qPCR) assays are provided in Supplementary Table 2. Technical replicate Cq values were averaged before calculation of relative transcript abundance. The number of independent biological replicates is reported in the corresponding figure legends.

For analysis of NMD-associated transcripts in H838 cells, 1 µg total RNA was reverse-transcribed using random hexamer primers and AffinityScript Multi-Temperature Reverse Transcriptase (Agilent Technologies, Cat. No. 600109). Quantitative PCR was performed in technical duplicate using Brilliant III Ultra-Fast SYBR Green qPCR Master Mix (Agilent Technologies, Cat. No. 600883) on a Rotor-Gene Q real-time PCR instrument (Qiagen). Reactions were performed in a final volume of 15 µL and contained cDNA corresponding to 24 ng RNA input equivalent and 0.5 µM each forward and reverse primer. Expression of the NMD-sensitive transcripts *HNRNPL*-NMD, *RPSP*, and *TRA2B*-NMD and the non-NMD *TRA2B* transcript was normalized to *ACTB* and expressed relative to the corresponding DMSO vehicle control.

For analysis of interferon-stimulated gene expression in H838 cells, cDNA was independently synthesized from 1 µg of the same RNA preparations using the High-Capacity cDNA Reverse Transcription Kit (Applied Biosystems, Cat. No. 4368813). Quantitative PCR was performed in technical triplicate in a final reaction volume of 10 µL using GoTaq qPCR Master Mix (Promega, Cat. No. A6001) on a QuantStudio 3 Real-Time PCR System (Applied Biosystems). Reactions contained 0.5 µM each gene-specific primer and 1 µL of 1:5-diluted cDNA, corresponding to 10 ng RNA input equivalent. Expression of *IFI44L*, *ISG15*, and *MX1* was normalized to *ACTB* and expressed relative to the corresponding DMSO vehicle control.

For analysis of T-cell activation-associated gene expression following co-culture with non-cryopreserved CD8⁺ T-cells, RNA from parallel co-culture samples was reverse-transcribed using the PrimeScript RT Reagent Kit (Takara, Cat. No. RR037A). Quantitative PCR was performed in technical triplicate in a final reaction volume of 10 µL using PowerUp SYBR Green Master Mix (Applied Biosystems, Cat. No. A25743) on a QuantStudio 7 Flex Real-Time PCR System (Applied Biosystems). Reactions contained cDNA corresponding to 4.5 ng RNA input and 0.5 µM each forward and reverse primer. Expression of *IFNG*, *GZMB*, *PRF1*, *IL2* and *TNF* was normalized to *CD8A* and expressed relative to the unmatched-donor DMSO control.

### Long-Read RNA Sequencing and Isoform Analysis

H838 cells were treated with 0.6 µM SMG1i or 0.1% DMSO for 24 h, with three technical replicates per condition. Total RNA was isolated using the Maxwell RSC simplyRNA Cells Kit (Promega, Cat. No. AS1390). RNA quantity and quality were assessed using a Qubit 4.0 fluorometer with the Qubit RNA BR Assay Kit (Thermo Fisher Scientific, Cat. No. Q10211) and a Fragment Analyzer system with the RNA Kit (Agilent Technologies, Cat. No. DNF-471), respectively. Kinnex full-length RNA libraries were generated from 300 ng total RNA according to the PacBio Kinnex full-length RNA workflow. Barcoded cDNA libraries were pooled and processed for Kinnex library generation. Final libraries were quantified using the Qubit dsDNA HS Assay Kit (Thermo Fisher Scientific, Cat. No. Q32854), and fragment size was assessed using a FEMTO Pulse system with the Genomic DNA 165 kb Kit (Agilent Technologies, Cat. No. FP-1002-0275). Six libraries were sequenced across two Revio SMRT Cells on a PacBio Revio system. Library preparation, sequencing, and primary processing were performed by the Next Generation Sequencing Platform, University of Bern.

Differential isoform abundance between SMG1i- and DMSO-treated samples was assessed using the Bioconductor R package edgeR ^72^. The edgeR-derived log2 fold change (log₂FC) represented the estimated base-2 log fold difference in isoform abundance between SMG1i-treated and DMSO-treated samples, with positive values indicating increased abundance following SMG1 inhibition. Isoforms were classified as significantly increased or decreased using FDR < 0.05 and |log₂FC| > log₂(1.5). Strict NMD annotation was assigned at the transcript level based on a GENCODE v39 *nonsense_mediated_decay* biotype and/or high-confidence NMDtxDB evidence defined by the presence of a premature termination codon together with significant accumulation following perturbation of an NMD factor. ^38,39^. The combined strict class included isoforms supported by either annotation source as well as isoforms supported by both sources. For significantly increased strict NMD-annotated isoforms, the range and median of the edgeR-derived log₂FC values were calculated. Enrichment of strict NMD annotations among significantly increased isoforms was evaluated relative to all isoforms not classified as significantly increased using Fisher’s exact test. Enrichment was reported as an odds ratio with a two-sided 95% confidence interval.

### Ribosome Profiling and Data Processing

H838 cells were treated with 0.3 µM SMG1i or 0.1% DMSO for 24 h, with three technical replicates per condition. Cell lysates were digested with RNase I (Jena Bioscience, Cat. No. EN-176S) and monosomes were isolated by ultracentrifugation through 10-50% sucrose gradients. Ribosome-protected fragments were isolated and libraries prepared following established ribosome-profiling procedures^73–75^. Fragments of 28-32 nucleotides were size-selected, processed for library generation, and rRNA was depleted using custom biotinylated antisense oligonucleotides listed in Supplementary Table 3. Libraries were sequenced on an Illumina NextSeq 1000 using a P2 flow cell by the Next Generation Sequencing Platform, University of Bern.

### Cell-Specific H838 Protein Sequence Database Generation

PacBio Iso-Seq reads were merged across sequencing runs for each sample and processed using IsoSeq3 v4.3.0, pbmm2 v1.17.0, and pigeon v1.17.0. Reads were clustered, aligned to the GENCODE GRCh38.p14 reference genome (release 48)^76^, collapsed, classified, and filtered using the PacBio workflow. ribo-seq data were adapter-trimmed using cutadapt v2.6 and processed using riboseq-flow v1.1.1^77^, including UMI extraction and deduplication and removal of contaminating RNA reads.

For generation of the cell-specific protein sequence database used in this study, ribo-seq reads were mapped directly to the filtered Iso-Seq transcript sequences. Iso-Seq sequences were extended by 18 nucleotides at both termini using the corresponding genomic sequence, and ribo-seq reads were mapped using Bowtie v1.3.1, allowing a maximum of one mismatch and retaining best-stratum alignments. After UMI deduplication, translating open reading frames (ORFs) were identified using ribotricer v1.4.0^78^ with a phase-score cutoff of 0.440 and a minimum of five valid codons. ORFs detected in at least two replicates of either experimental group were retained. Identical translated protein sequences were collapsed, and sequences fully contained within longer protein sequences were removed, resulting in 398,930 H838-derived protein sequence entries.

### Whole Cell Proteome and Phosphoproteome Analysis

H838 cells were treated as described above and lysed in 8 M urea, 100 mM Tris-HCl (pH 8.0), 10 mM NaF, 2 mM Na₃VO₄, and protease inhibitor cocktail (Roche, Cat. No. 11873580001). Lysates were sonicated, clarified by centrifugation, and protein concentrations were determined by Bradford assay. Proteins were reduced with 10 mM dithiothreitol, alkylated with 50 mM iodoacetamide, and precipitated with ice-cold acetone. For digestion, 640 µg protein in 8 M urea was diluted to 1.6 M with four volumes of 20 mM Tris-HCl containing 2 mM CaCl₂. Trypsin (5 µg; protein-to-enzyme ratio 128:1, w/w) was added for 2 h at 37 °C, followed by a second 5 µg addition and overnight digestion at room temperature, resulting in a cumulative ratio of 64:1. Digestion was stopped with 1% TFA, and samples were clarified at 16,000 × *g* for 10 min. Peptides were desalted using Oasis C18 cartridges (Waters, Cat. No. WAT054955) and phosphopeptides were enriched on a Bravo AssayMAP platform using Fe-NTA cartridges (Agilent Technologies, Cat. No. G5496-60085). Phosphopeptides were eluted with 1% NH₄OH, immediately acidified, and analyzed by nLC-MS/MS).

Mass-spectrometric analysis was performed on an Ultimate 3000 system (Thermo Fisher Scientific) coupled to a timsTOF Pro mass spectrometer (Bruker Daltonics). Peptides were separated on a custom-packed C18 CSH analytical column (75 µm × 20 cm, 1.7 µm) at 250 nL/min using a 70-min gradient from 2% to 40% ACN in 0.1% formic acid. Quantitative phosphoproteomic data were acquired by DIA-PASEF using 32 precursor isolation windows covering 400-1,200 *m/z* together with a corresponding DDA-PASEF acquisition using the same parameters like with the first set of immunopeptidomics analysis (see below).

Mass-spectrometry data were interpreted using Spectronaut version 21.0.260602.94842 (Biognosys AG) with the BGS Phospho PTM Workflow in hybrid directDIA mode where the DDA runs were used to support spectra library building. Data interpretation was done with the same protein sequence database as for the immunopeptidomics data. Carbamidomethylation of cysteine was set as a fixed modification, whereas protein N-terminal acetylation, methionine oxidation, and phosphorylation of serine, threonine, and tyrosine were set as variable modifications, allowing up to five variable modifications per peptide. Identifications were controlled at 1% false-discovery rate, and only phosphosites with a localization probability ≥0.75 were retained. Quantification was based on MS2 peak areas, and automatic cross-run normalization was applied in Spectronaut.

### Production and purification of W6/32 monoclonal antibody

W6/32-producing HB-95 hybridoma cells (ATCC, HB-95) were transferred to a CELLine CL 350 two-compartment bioreactor flask (INTEGRA Biosciences) and maintained according to the manufacturer’s instructions. Briefly, the cell compartment was filled with complete medium, while the nutrient reservoir was filled with DMEM containing 0.8% FBS and 1% P/S. After approximately 7-8 days, when viability had decreased to approximately 50%, the antibody-containing medium was collected from the cell compartment.

W6/32 antibodies were purified by gravity-flow chromatography using 1 mL Protein A-Sepharose 4B resin (ThemoFisher Scientific, Cat. No. 101042). The resin was equilibrated with 10 column volumes (c.v.) 0.1 M Tris-HCl, pH 8.0, before the clarified culture supernatant was loaded. After washing, bound antibodies were eluted with 0.1 N acetic acid, pH 3.0, directly into a neutralization buffer with final concentrations of 100 mM boric acid, 75 mM NaCl, and 25 mM sodium tetraborate. The pH was subsequently adjusted to pH 8. Antibody concentrations were determined and aliquots were stored at −20 °C until use.

### W6/32 affinity column preparation

A 500-µL Protein-A Sepharose 4B resin bed (Invitrogen, Cat. No. 101042) was prepared in a Bio-Rad Poly-Prep chromatography column (Bio-Rad). Purified W6/32 antibody (10-12 mg) was loaded onto the resin and incubated for 1 h at room temperature under end-over-end rotation.

The antibody-bound resin was washed twice with 10 column volumes of 0.2 M boric acid (pH 9.0). Cross-linking was performed with four column volumes of 40 mM dimethyl pimelimidate dihydrochloride (DMP, Thermo Scientific, Cat. No. 21667), freshly dissolved in 0.2 M boric acid (pH 9.0), for 30 min at room temperature under end-over-end rotation. The reaction was quenched with nine column volumes of 0.2 M Tris-HCl (pH 8.0) for 30 min at room temperature. The resin was then washed sequentially with 20 column volumes of 1% acetic acid and 10 column volumes of Tris-buffered saline (50 mM Tris-HCl, pH 7.6, 150 mM NaCl, 0.05% NaN₃). The coupled resin was stored as a 1:1 slurry in PBS containing 0.02% NaN₃ at 4 °C. After use, columns were regenerated with 1% acetic acid, re-equilibrated in PBS containing 0.02% NaN₃, and reused for subsequent experiments up to fifteen times.

### Peptide-HLA-I complex enrichment

Frozen H838 cell pellets corresponding to approximately 1 × 10⁷ cells were lysed in 1 mL ice-cold buffer containing 0.5% NP-40 Alternative (Calbiochem, Cat. No. 492016), 50 mM Tris-HCl (pH 8.0), 150 mM NaCl, and EDTA-free protease inhibitor cocktail (Roche, Cat. No. 11873580001).

Clarified lysates were pre-cleared using Protein A-Sepharose 4B (Invitrogen, Cat. No. 101042) and incubated with 500 µL W6/32-coupled affinity resin for 1 h at 4 °C under end-over-end rotation. The resin was washed sequentially with 10 c.v. each of WB1 (0.005% IGEPAL CA-630, 50 mM Tris-HCl, pH 8.0, 150 mM NaCl, 5 mM EDTA, 100 µM PMSF, and 1 µg/mL pepstatin A), WB2 (50 mM Tris-HCl, pH 8.0, 150 mM NaCl), WB3 (50 mM Tris-HCl, pH 8.0, 450 mM NaCl), and WB4 (50 mM Tris-HCl, pH 8.0). Peptide-HLA-I complexes were eluted with 1% acetic acid and lyophilized. Prior to C18 purification, lyophilized material was resuspended in 120 µL 20% formic acid.

### C18 solid-phase extraction of HLA-I peptides

HLA eluates were subjected to C18 solid-phase extraction using three plugs punched from a 3M Empore C18 extraction disk (Sigma-Aldrich, Cat. No. 66883-U) packed into a 200-µL pipette tip. The C18 material was activated with methanol and equilibrated with 0.1% trifluoroacetic acid (TFA), with centrifugation at 3,000 × g between steps.

Samples were loaded onto the C18 material and washed with 100 µL 0.1% TFA. Bound peptides were eluted sequentially with 40 µL of 20% acetonitrile (ACN)/0.1% TFA (Fraction 1), followed by 40 µL of 40% ACN/0.1% TFA (Fraction 2). Fractions were dried under a gentle stream of nitrogen and reconstituted in 20 µL 20% formic acid before nLC-MS/MS analysis.

### Immunopeptidomics nLC-MS/MS Analysis

Two batches of immunopeptide replicates were analyzed on the same LC-MS/MS instrument as described above using two different LC gradient and MS methods. After loading, peptides were eluted in back flush mode onto a homemade C18 CSH Waters column (1.7 μm, 130 Å, 75 μm × 20 cm) by applying a biphasic 70- or 50-minute acetonitrile gradient, for a first and a second batch of samples, respectively. The acetonitrile concentration went from 2% to 35% in water / 0.1% formic acid, at a flow rate of 250 nL/min. The timsTOF Pro was operated in the parallel accumulation-serial fragmentation (PASEF) mode. The mass range was set between 100 and 1700 m/z, with 10 or 6 PASEF scans between 0.7 and 1.76 or 0.6 and 1.70 V s/cm^2^, an accumulation time of 2 ms or 200 ms, and the ramp time of 166 ms or 200 ms, respectively. Fragmentation was triggered at 14,000 arbitrary units (au), and peptides (up to charge 5 or 3) were fragmented using collision induced dissociation with a spread between 20 to 70 or 20 to 60 eV. Mass spectrometry data was then interpreted with FragPipe v24^79^ against the final H838-specific proteogenomic sequence database generated from PacBio Kinnex long-read RNA-sequencing and ribosome-profiling data, concatenated with the human reference proteome downloaded from UniProtKB (version 2025_12), and with common mass-spectrometry contaminant proteins. An unspecific digestion setting was used for the database search. The following modifications were set as variable: protein N-terminal acetylation, cysteinylation of cysteine, oxidation of methionine, pyro-glu from glutamine and asparagine. Only one such modification was allowed per peptide, precursor and fragment mass error tolerances were set at ±20 ppm and ±0.04 Da, and peptides and peptide spectrum matches were filtered to a 1% FDR based on a concatenated target and decoy search, respectively.

### Extended Metascore Calculation

Peptide identifications were used as input for the extended-metascore workflow. Candidate peptides were evaluated for predicted binding using NetMHCpan 4.2^36^. The prediction panel included the three genotyped H838 HLA-I alleles (HLA-A*30:02, HLA-B*18:01, HLA-C*05:01) plus three additional alleles (HLA-A*32:01, HLA-A*74:01, HLA-B*48:01). Binder classifications were assigned using the NetMHCpan default percentile-rank thresholds for strong, weak, and non-binders.

For each peptide-HLA-I combination, the metascore integrated the NetMHCpan-predicted affinity (Aff), eluted-ligand score (ELscore), binding-affinity score (BAscore), peptide-length term (*L*_pep_), and binder-level term (*BL*_pep_):

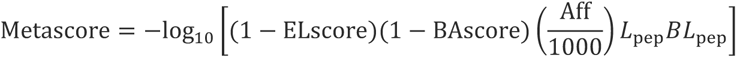

Aff represents the predicted peptide-HLA binding affinity in nM. *L*_pep_ was set to 1 for peptides shorter than 15 amino acids and to 5 for longer peptides. *BL*_pep_ was set to 1 for strong binders, 2 for weak binders, and 10 for non-binders.

Proteasomal cleavage predictions were generated using NetChop 3.1 with the C-terminal and 20S models^37,80^. Cleavage sites with scores of at least 0.5 were retained. The model-specific outputs were merged with the peptide-level NetMHCpan results, yielding separate N- and C-terminal cleavage scores for each NetChop model. Missing cleavage scores were annotated as zero.

When evidence of cleavage was present at both the N- and C-terminal peptide ends, the maximum cleavage score was added to the metascore. When cleavage evidence was present at only one peptide end, half of the maximum score was added. In the absence of predicted cleavage at either peptide end, the original metascore was retained. For peptides evaluated against multiple HLA-I alleles, the allele yielding the highest extended metascore was retained.

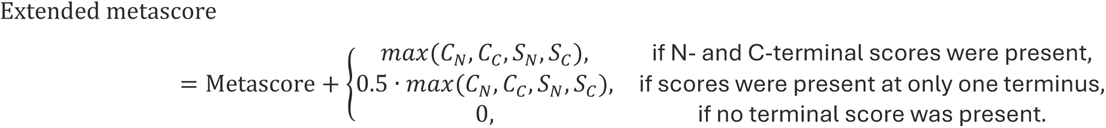

The extended-metascores formed a bimodal, nearly Gaussian distribution from where a 1% peptide-level FDR could be calculated using the target-decoy approach (FDR = number of decoy hits above threshold / number of target hits above threshold). This threshold was subsequently applied during peptide filtering (see “Immunopeptidomics Data Evaluation”). All computational analyses were performed in R version 4.5.2 (R Core Team, 2025).

### Immunopeptidomics Data Evaluation

Peptide entries retained the sequence and protein annotations encoded in the cell-specific database, including appended UniProtKB annotations where available. Peptides classified as passing the 1% peptide-level FDR in the extended-metascore output were retained for subsequent evaluation.

For treatment classification, peptide-spectrum matches (PSMs) were summed for each peptide across SMG1i-treated and DMSO-treated samples. Peptides with fewer than four PSMs in both conditions were considered insufficiently supported. Entries with 0 DMSO PSM and ≥4 SMG1i PSM were classified as SMG1i-only, whereas entries with ≥4 DMSO PSM and 0 SMG1i PSM were classified as DMSO-only. For entries detected in both conditions, the SMG1i/DMSO PSM ratio was used to classify peptides as SMG1i-enriched (ratio >2), DMSO-enriched (ratio <0.5), or otherwise unchanged. Based on this classification, SMG1i-only candidates were further prioritized after exclusion of contaminant-derived entries. Candidates were required to be unmodified 9-to 11-mer peptides, detected in at least three SMG1i-treated samples, and predicted to bind one of the genotyped H838 HLA-I alleles (HLA-A*30:02, HLA-B*18:01, or HLA-C*05:01).

For motif analysis, sequence logos were generated from unique 9-mer peptide sequences assigned to selected HLA alleles using the ggseqlogo R package_81_ in R version 4.5.2 (R Core Team, 2025), with information-content scaling in bits and without PSM weighting.

### Immunofluorescence staining

H838 cells were grown in eight-well chamber slides (Ibidi, Cat. No. 81201) and treated for 24 h with 0.2 or 0.3 µM SMG1i or 0.1% DMSO vehicle control. Staining was performed without permeabilization using an Alexa Fluor 647-conjugated W6/32 antibody (Santa Cruz Biotechnology, Cat. No. SC-32235), a conformation-dependent pan-HLA-I antibody recognizing surface-expressed, properly folded HLA-I complexes, at a 1:100 or 1:300 dilution, or an Alexa Fluor 647-conjugated mouse IgG2a isotype control (Santa Cruz Biotechnology, Cat. No. SC-24637) at the corresponding dilution.

Images were acquired using an inverted Carl Zeiss LSM 980 laser-scanning confocal microscope equipped with a Plan-Apochromat 63×/1.4 oil-immersion objective at the Live Cell Imaging Core Facility, University of Bern.

### Flow Cytometric Analysis of Surface HLA-I

H838 cells were treated for 24 h with 0.2 or 0.3 µM SMG1i or 0.1% DMSO vehicle control and detached using 4 mM EDTA in DPBS. Cells were resuspended in FACS buffer consisting of DPBS supplemented with 5% FBS.

Fc receptors were blocked with Human TruStain FcX (BioLegend, Cat. No. 422301) for 10 min on ice. Surface HLA-I was stained for 30 min on ice in the dark using Alexa Fluor 647-conjugated W6/32 pan-HLA-I antibody (Santa Cruz Biotechnology, Cat. No. sc-32235) at 1:640 or 1:1280 dilution. Alexa Fluor 647-conjugated mouse IgG2a (Santa Cruz Biotechnology, Cat. No. sc-24637) was used as the corresponding isotype control.

DAPI was added immediately before acquisition. Data were acquired on a CytoFLEX S flow cytometer (Beckman Coulter), recording 50,000 events per sample. Debris, doublets, and DAPI-positive cells were excluded by sequential gating (Supplementary Figure 4), and surface HLA-I abundance was quantified as the median AF647-A fluorescence intensity of viable singlet cells. Three technical staining and acquisition replicates were analyzed per condition.

### CD8⁺ T-cell isolation and activation

PBMCs were obtained from three HLA-typed healthy donors, including two donors sharing 3/6 HLA-A, HLA-B, and HLA-C alleles with H838 and one unmatched donor (0/6). PBMCs were isolated from fresh whole blood using BD Vacutainer CPT tubes (BD Biosciences, Cat. No. 362782) and either used immediately or cryopreserved in 90% FBS and 10% DMSO and stored in liquid nitrogen.

CD8⁺ T-cells were isolated by negative selection using the EasySep Human CD8⁺ T Cell Isolation Kit (STEMCELL Technologies, Cat. No. 17913). CD8⁺ T cell purity was assessed by flow cytometry on a FACSAria III cell sorter (BD Biosciences) using FITC anti-human CD3 (BioLegend, 317305), PE anti-human CD8 (BioLegend, 344705), and Alexa Fluor 700 anti-human CD45 (BioLegend, 368513) antibodies with DAPI viability exclusion. Populations with at least 90% of DAPI^-^CD45^+^CD3^+^CD8⁺ isolated cells were considered sufficiently pure for the experiments (Supplementary Figure 3).

Isolated CD8⁺ T-cells were maintained in DMEM medium supplemented with 2% FBS and 1× insulin-transferrin-selenium (ITS, Gibco, Cat. No. 41400045) and activated with Dynabeads Human T-Cell Activator CD3/CD28 (Thermo Fisher Scientific, Cat. No. 11161D) at a 1:1 bead-to-cell ratio for 10 h for freshly isolated cells or 24 h for cells isolated from cryopreserved PBMCs. Beads were removed by magnetic separation before co-culture.

### H838-CD8⁺ T-cell co-culture and intracellular cytokine staining

H838 cells were treated with 0.3 µM SMG1 inhibitor or 0.1% DMSO for 24 h before addition of 6 × 10⁴ activated CD8⁺ T-cells per well at an effector-to-target ratio of 2:1. Cells were co-cultured for 8 h. Brefeldin A (Sigma, Cat. No. B6542) was added 2 h after initiation of the co-culture at 5 µg/mL. CD8⁺ T-cell monocultures served as negative controls, whereas monocultures stimulated with 1× Cell Activation Cocktail without Brefeldin A (BioLegend, Cat. No. 423301) served as positive controls. Brefeldin A was added after 1 h of control stimulation and remained present for the final 4 h.

Following co-culture, nonadherent cells were collected and adherent cells were detached with 4 mM EDTA in DPBS and combined with the corresponding nonadherent fraction. Cells were stained with eBioscience Fixable Viability Dye eFluor 780 (Thermo Fisher Scientific, Cat. No. 65-0865-14), FITC anti-human CD3 (clone OKT3, BioLegend, Cat. No. 300406), and APC anti-human CD8 (clone RPA-T8, BioLegend, Cat. No. 301014). Following fixation and permeabilization using the eBioscience Foxp3/Transcription Factor Staining Buffer Set (Thermo Fisher Scientific, Cat. No. 00-5523-00), intracellular cytokines were detected using BV421 anti-human IFN-γ (clone B27, BioLegend, Cat. No. 506538) and PE anti-human IL-2 (clone MQ1-17H12, BioLegend, Cat. No. 500306).

Data were acquired on a BD LSRFortessa flow cytometer and analyzed using BD FACSDiva and FlowJo v10.10.1. T-cell responses were quantified as the percentage of IFN-γ⁺ or IL-2⁺ cells within viable CD3⁺CD8⁺ singlets. For experiments using CD8⁺ T-cells isolated from non-cryopreserved PBMCs, parallel co-cultures were prepared for analysis of T-cell activation-associated gene expression as described below.

### Experimental Design and Statistical Rationale

Immunopeptidomics comprised 0.2 µM SMG1i for 24 h (n = 3), 0.3 µM for 24 h (n = 3), 0.3 µM for 8 h (n = 3), and 0.6 µM for 24 h (n = 6), with equal numbers of corresponding DMSO controls. CD8⁺ T-cell co-culture experiments were performed in duplicate for three donors using cryopreserved PBMCs and in triplicate for two donors using non-cryopreserved PBMCs. Statistical tests, multiple-comparison procedures, replicate definitions, and data presentation are specified in the corresponding Methods and figure legends. No randomization or blinding was applied.

## Data and Code Availability

The mass spectrometry proteomics data have been deposited to the ProteomeXchange Consortium via the PRIDE^82^ partner repository with the dataset identifier PXD087616. The PacBio Kinnex long-read RNA sequencing and Illumina ribosome profiling (Ribo-seq) data have been deposited in the NCBI Gene Expression Omnibus (GEO)^83,84^ under accession numbers GSE354640 and GSE355017, respectively.

## Ethics statement

The Cantonal Ethics Committee of Bern determined that the project did not fall within the scope of the Swiss Human Research Act and therefore did not require ethics committee authorization (Req-2026-01151).

## Acknowledgements

We first thank Prof. Daniel Sidler, Inselspital Bern, for facilitating access to donor-derived material and for essential experimental and logistical support in establishing the PBMC and CD8⁺ T-cell experiments. His support was central to implementing the functional immune-recognition component of this study. We thank Prof. Thomas Marti and his laboratory at the Department for BioMedical Research (DBMR), University of Bern, for kindly providing the H838 cell line and for access to laboratory resources used for interferon-stimulated gene expression analysis. We further thank Christel Dubey for technical guidance. We acknowledge the Next Generation Sequencing Platform and Pamela Nicholson for sequencing support. We thank Daniel Schärer and Michael Horn at the Zentrum für Labormedizin, Inselspital, for support with HLA genotyping. We acknowledge Miriam Diaz Varela and Thomas Schaffer of the DBMR Flow Cytometry and Cell Sorting Core Facility for technical support with flow-cytometric analyses, and Ekaterina Ivanova and the Microscopy Imaging Center for support with fluorescence microscopy. We further thank Prof. Sebastian Leidel and his group, Department of Chemistry, Biochemistry and Pharmaceutical Sciences, University of Bern, for providing reagents and laboratory support for the ribosome-profiling experiments. Finally, we gratefully acknowledge the anonymous blood donors whose samples contributed to this study.

## Author Contributions

A.B.: Conceptualization, Methodology, Validation, Formal analysis, Investigation, Data curation, Visualization, Project administration, Writing original draft, Writing review C editing. A.-C.U.: Methodology, Software, Formal analysis, Data curation, Writing review C editing. S.B.L.: Methodology, Investigation, Resources, Writing review C editing. M.B.: Methodology, Investigation, Writing review C editing. J.M.: Methodology, Investigation, Resources, Writing review C editing. O.M.: Conceptualization, Supervision, Resources, Funding acquisition, Writing review C editing. M.H.: Conceptualization, Methodology, Formal analysis, Resources, Supervision, Project administration, Funding acquisition, Writing review C editing.

## Supplementary Figures & Tables

**Supplementary Figure 1.**
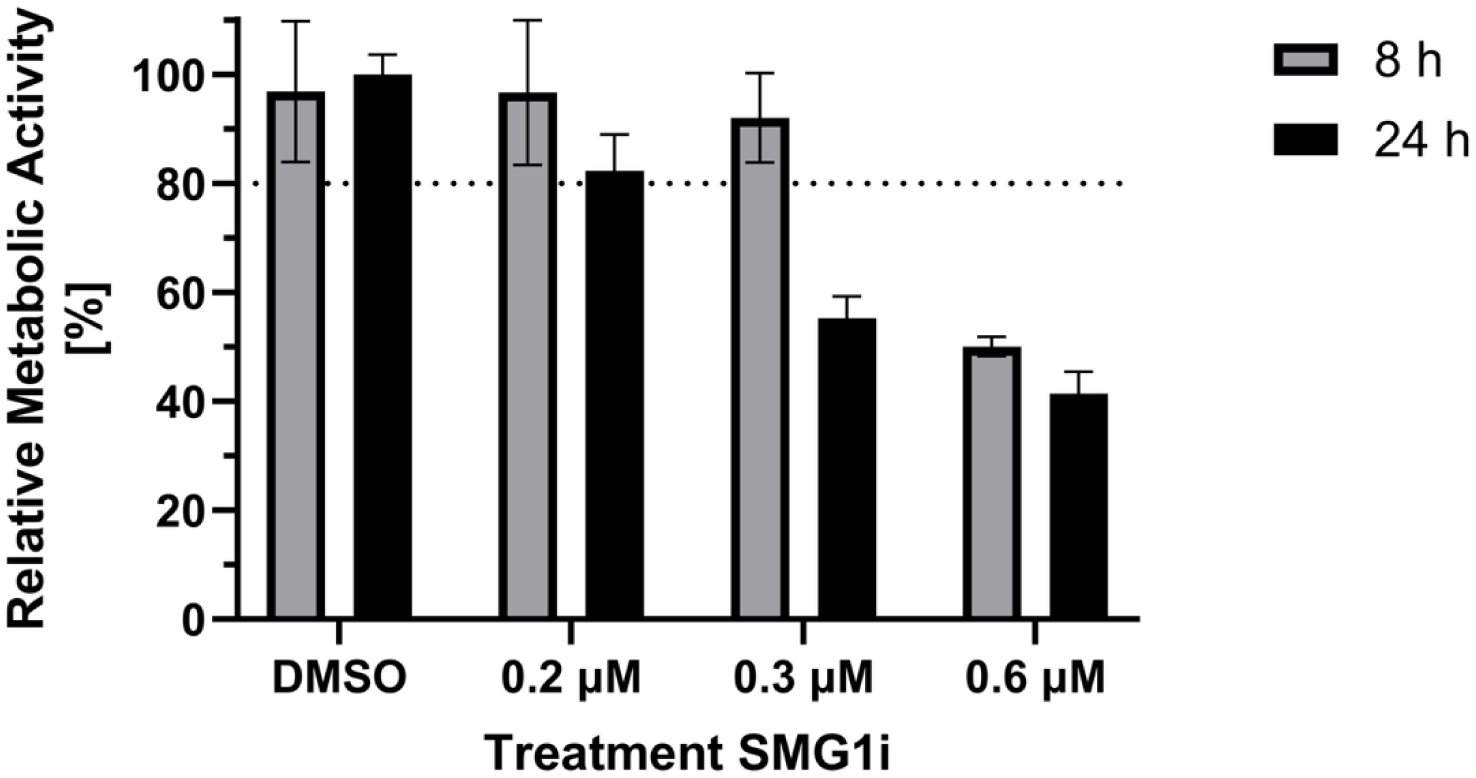
SMG1i concentration-dependent reduction in metabolic activity of H838 cells. H838 cells were treated with 0.1% DMSO vehicle control or increasing SMG1i concentrations (0.2-0.6 µM) for 8h or 24 h. Metabolic activity was assessed using an MTT assay and normalized to the vehicle control of 24 h treatment, which was set to 100%. Bars represent the mean ± SD of three plate-level technical replicates. Each plate-level value represents the mean of six replicate wells measured on a 96-well plate.

**Supplementary Figure 2.**
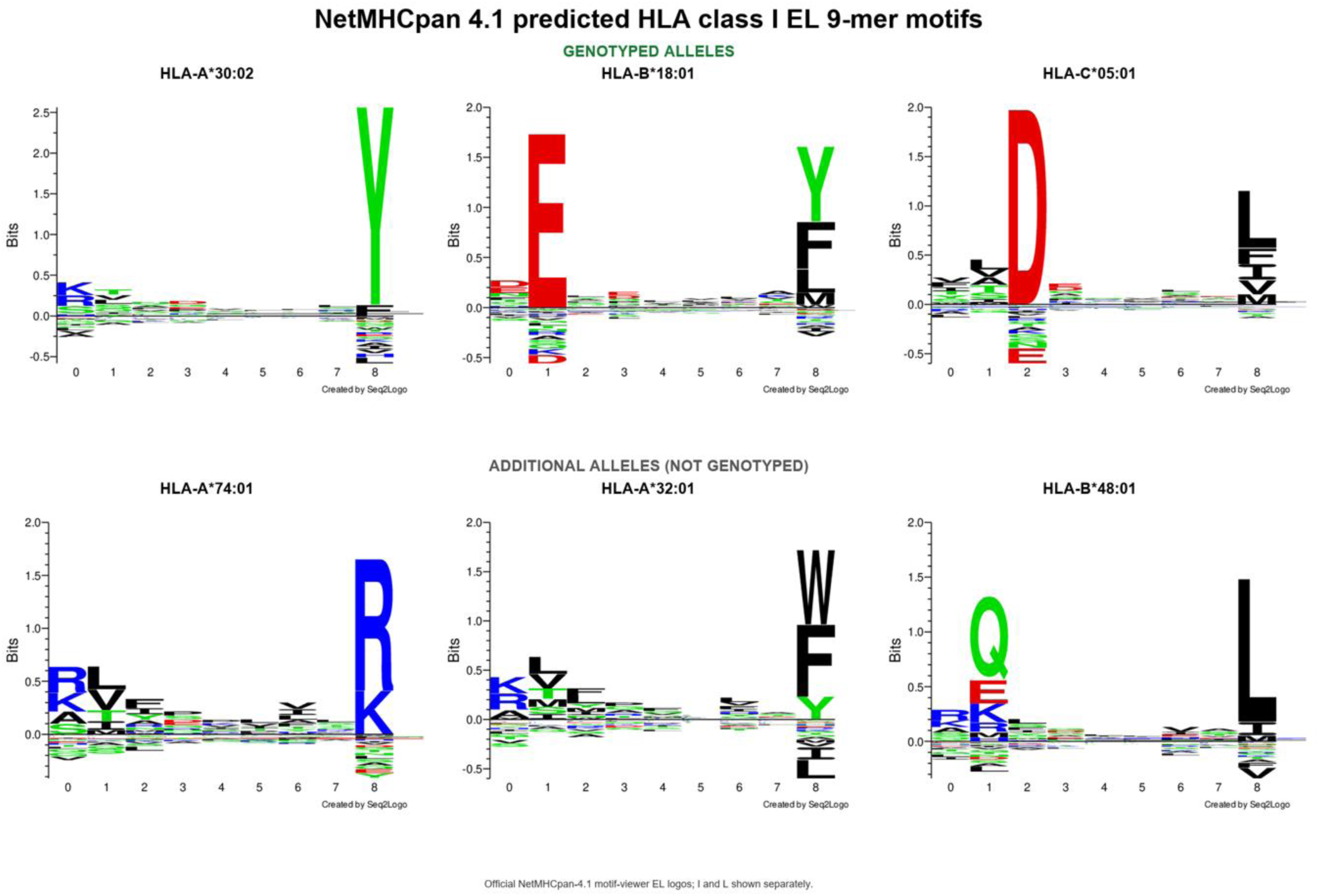
NetMHCpan-predicted HLA class I peptide-binding motifs. Sequence logos show predicted 9-mer binding motifs for the three genotyped H838 HLA alleles (HLA-A*30:02, HLA-B*18:01, HLA-C*05:01; upper row) and three additional alleles included in the prediction panel (HLA-A*74:01, HLA-A*32:01, HLA-B*48:01; lower row). HLA-B*48:01 has previously been reported for NCI-H838 and is listed in Cellosaurus, but was not detected by either sequencing-based HLA-genotyping approach in the H838 cells analyzed in this study. Motifs were obtained from the NetMHCpan 4.1 motif viewer. Amino acid preferences at each position are shown as information content (bits).

**Supplementary Figure 3.**
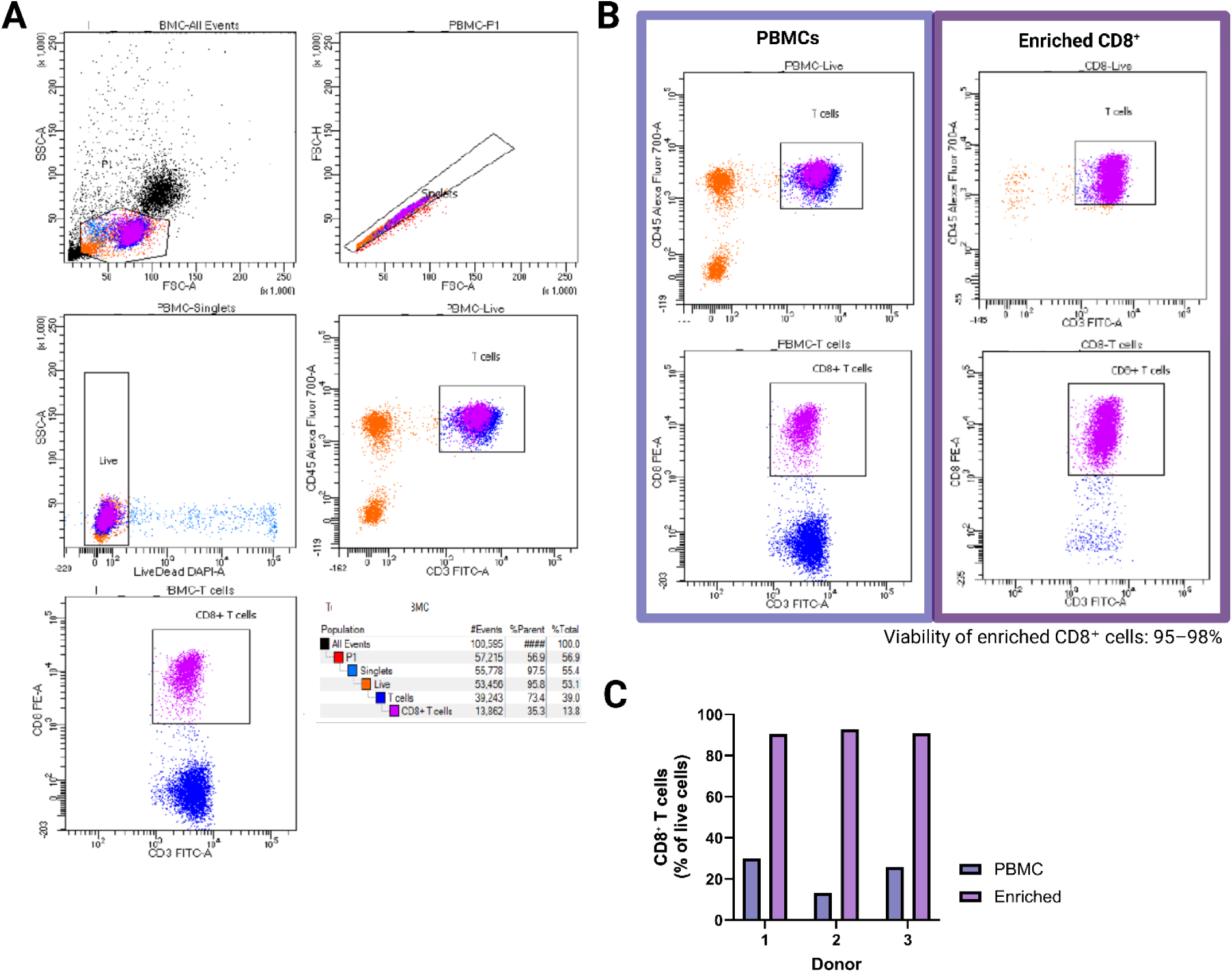
Flow-cytometric validation of CD8⁺ T-cell enrichment from PBMCs. **(A)** Representative flow-cytometric gating strategy used to assess CD8⁺ T-cell purity. Following exclusion of debris and doublets, viable cells were selected by DAPI exclusion. T cells were identified as CD45⁺CD3⁺ cells, and CD8⁺ T cells were subsequently identified based on CD8 expression. **(B)** Representative comparison of peripheral blood mononuclear cells (PBMCs) before enrichment and the corresponding CD8⁺ T-cell-enriched fraction following negative selection using the EasySep Human CD8⁺ T Cell Isolation Kit. CD45/CD3 and CD3/CD8 staining demonstrates enrichment of the CD45⁺CD3⁺CD8⁺ T-cell population. Viability of the enriched fractions was 95–98%. **(C)** CD8⁺ T cells as a percentage of viable cells before and after enrichment for three representative donor preparations, showing enrichment to approximately 90% or greater CD8⁺ T-cell purity.

**Supplementary Figure 4.**
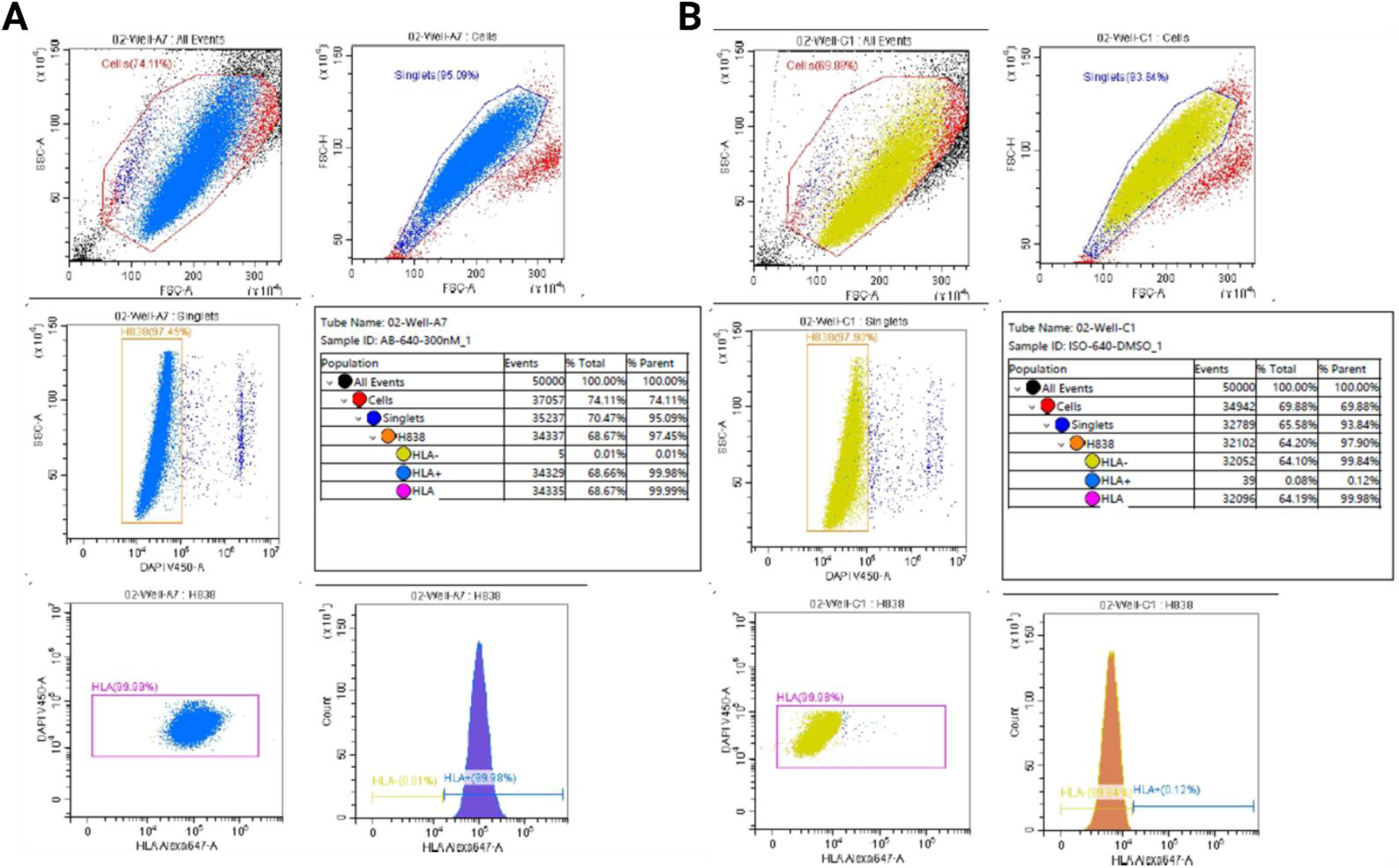
Flow-cytometric assessment of HLA-I surface expression on H838 cells. Representative flow-cytometric gating strategy used to quantify surface HLA-I expression on H838 cells. Debris and doublets were excluded based on forward- and side-scatter characteristics, followed by exclusion of DAPI-positive cells to retain viable singlets. (**A)** Representative H838 sample stained with Alexa Fluor 647-conjugated W6/32, a conformation-dependent pan-HLA-I antibody recognizing properly folded cell-surface HLA-I complexes. **(B)** Corresponding representative sample stained with Alexa Fluor 647-conjugated mouse IgG2a isotype control.

**Supplementary Table 1.**
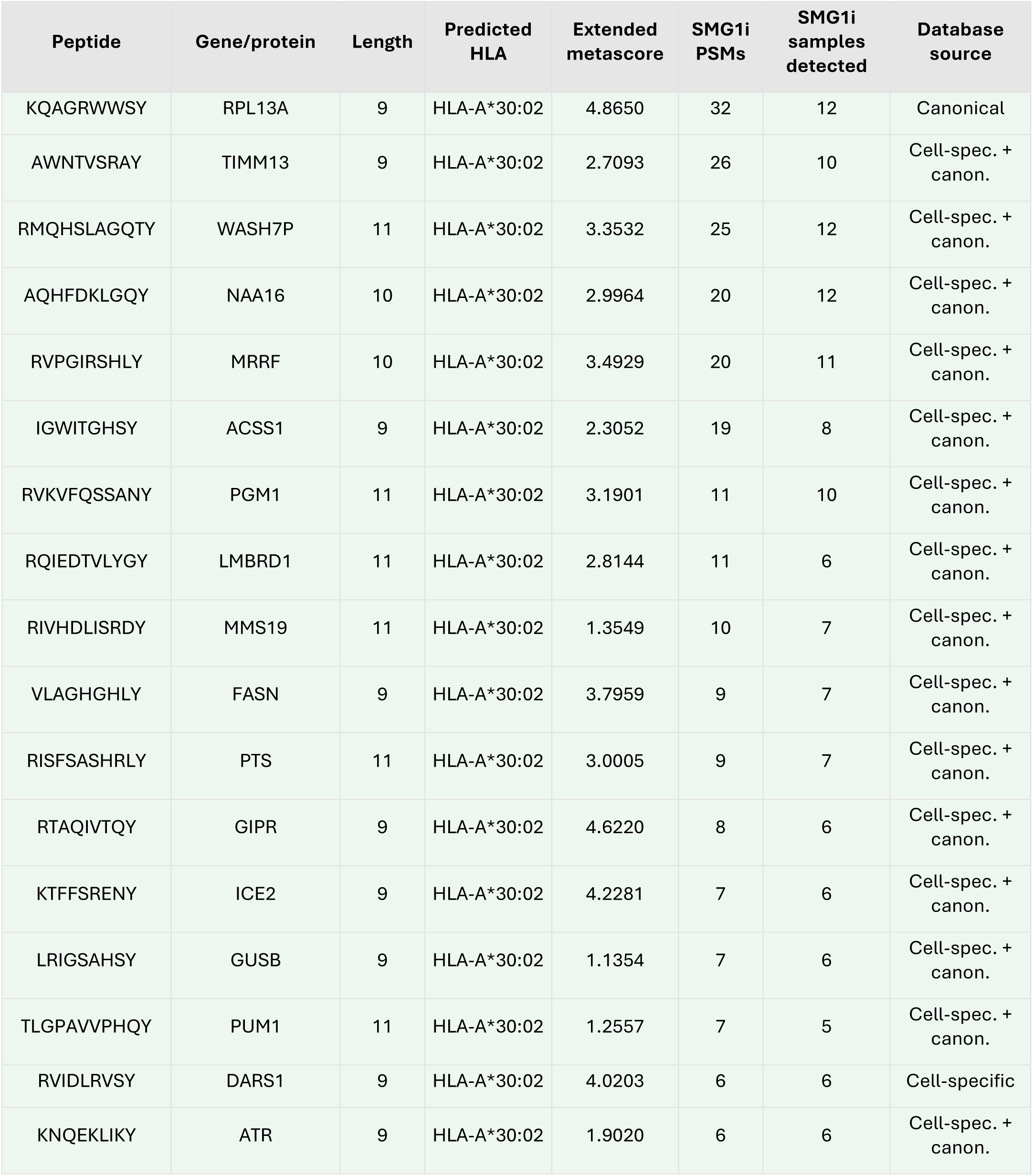
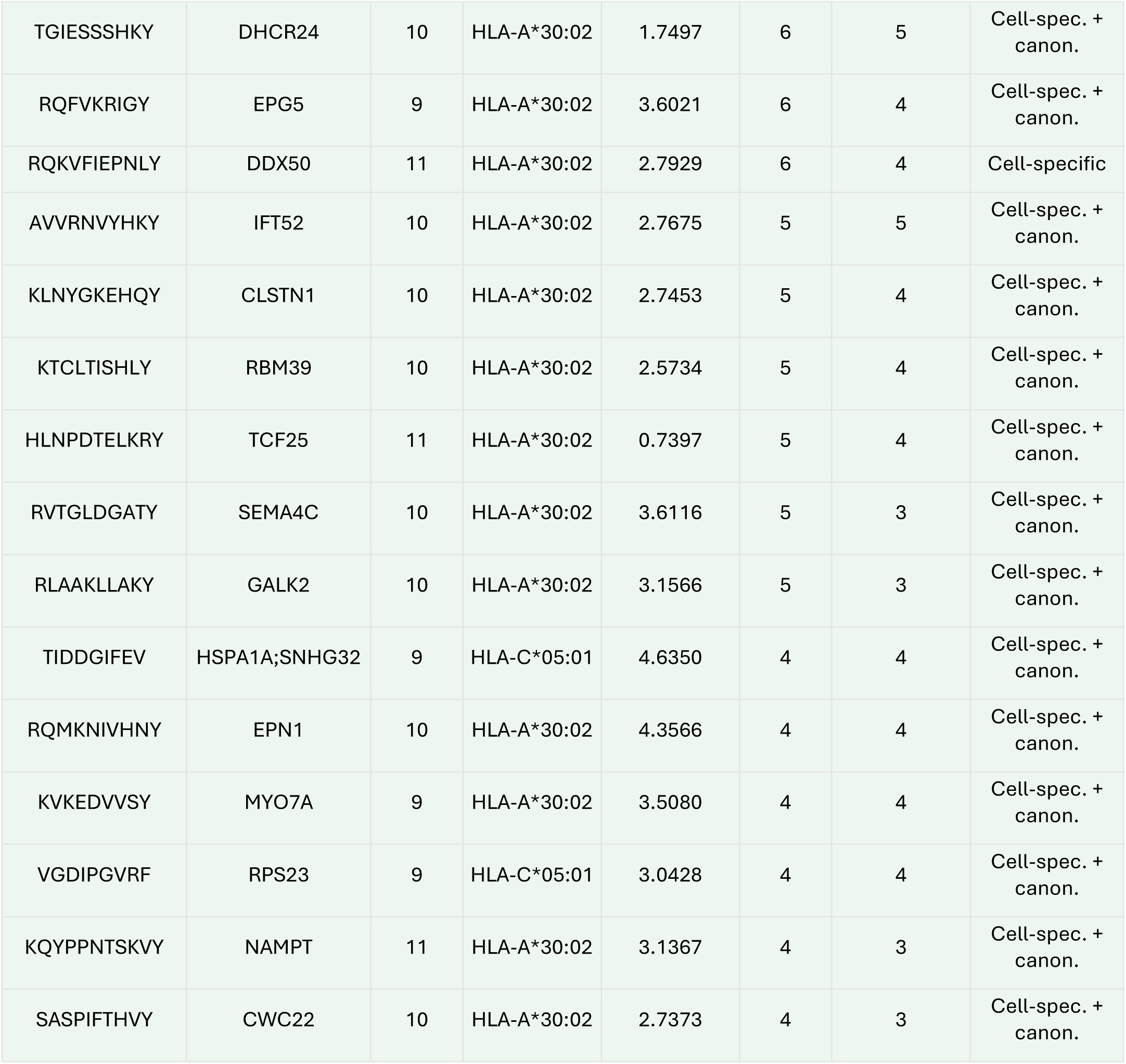
Prioritization of SMG1i-associated HLA-I peptide candidates identified in H838 cells. The table lists the 32 peptides retained after application of the final candidate-selection criteria described in the Methods section “Immunopeptidomics Data Evaluation.” For each peptide, the sequence, assigned gene/protein, peptide length, predicted HLA-I allele, extended metascore, number of SMG1i-associated peptide-spectrum matches (PSMs), number of SMG1i-treated samples in which the peptide was detected, and database source are reported. Abbreviations: PSM, peptide-spectrum match; SMG1i, SMG1 inhibitor; Cell-spec., cell-specific database; canon., canonical database.

| Peptide | Gene/protein | Length | Predicted HLA | Extended metascore | SMG1i PSMs | SMG1i samples detected | Database source |
|---|---|---|---|---|---|---|---|
| KQAGRWWSY | RPL13A | 9 | HLA-A*30:02 | 4.8650 | 32 | 12 | Canonical |
| AWNTVSRAY | TIMM13 | 9 | HLA-A*30:02 | 2.7093 | 26 | 10 | Cell-spec. + canon. |
| RMQHSLAGQTY | WASH7P | 11 | HLA-A*30:02 | 3.3532 | 25 | 12 | Cell-spec. + canon. |
| AQHFDKLGQY | NAA16 | 10 | HLA-A*30:02 | 2.9964 | 20 | 12 | Cell-spec. + canon. |
| RVPGIRSHLY | MRRF | 10 | HLA-A*30:02 | 3.4929 | 20 | 11 | Cell-spec. + canon. |
| IGWITGHSY | ACSS1 | 9 | HLA-A*30:02 | 2.3052 | 19 | 8 | Cell-spec. + canon. |
| RVKVFQSSANY | PGM1 | 11 | HLA-A*30:02 | 3.1901 | 11 | 10 | Cell-spec. + canon. |
| RQIEDTVLYGY | LMBRD1 | 11 | HLA-A*30:02 | 2.8144 | 11 | 6 | Cell-spec. + canon. |
| RIVHDLISRDY | MMS19 | 11 | HLA-A*30:02 | 1.3549 | 10 | 7 | Cell-spec. + canon. |
| VLAGHGHLY | FASN | 9 | HLA-A*30:02 | 3.7959 | 9 | 7 | Cell-spec. + canon. |
| RISFSASHRLY | PTS | 11 | HLA-A*30:02 | 3.0005 | 9 | 7 | Cell-spec. + canon. |
| RTAQIVTQY | GIPR | 9 | HLA-A*30:02 | 4.6220 | 8 | 6 | Cell-spec. + canon. |
| KTFFSRENY | ICE2 | 9 | HLA-A*30:02 | 4.2281 | 7 | 6 | Cell-spec. + canon. |
| LRIGSAHSY | GUSB | 9 | HLA-A*30:02 | 1.1354 | 7 | 6 | Cell-spec. + canon. |
| TLGPAVVPHQY | PUM1 | 11 | HLA-A*30:02 | 1.2557 | 7 | 5 | Cell-spec. + canon. |
| RVIDLRVSY | DARS1 | 9 | HLA-A*30:02 | 4.0203 | 6 | 6 | Cell-specific |
| KNQEKLIKY | ATR | 9 | HLA-A*30:02 | 1.9020 | 6 | 6 | Cell-spec. + canon. |
| TGIESSSHKY | DHCR24 | 10 | HLA-A*30:02 | 1.7497 | 6 | 5 | Cell-spec. + canon. |
| RQFVKRIGY | EPG5 | 9 | HLA-A*30:02 | 3.6021 | 6 | 4 | Cell-spec. + canon. |
| RQKVFIEPNLY | DDX50 | 11 | HLA-A*30:02 | 2.7929 | 6 | 4 | Cell-specific |
| AVVRNVYHKY | IFT52 | 10 | HLA-A*30:02 | 2.7675 | 5 | 5 | Cell-spec. + canon. |
| KLNYGKEHQY | CLSTN1 | 10 | HLA-A*30:02 | 2.7453 | 5 | 4 | Cell-spec. + canon. |
| KTCLTISHLY | RBM39 | 10 | HLA-A*30:02 | 2.5734 | 5 | 4 | Cell-spec. + canon. |
| HLNPDTELKRY | TCF25 | 11 | HLA-A*30:02 | 0.7397 | 5 | 4 | Cell-spec. + canon. |
| RVTGLDGATY | SEMA4C | 10 | HLA-A*30:02 | 3.6116 | 5 | 3 | Cell-spec. + canon. |
| RLAAKLLAKY | GALK2 | 10 | HLA-A*30:02 | 3.1566 | 5 | 3 | Cell-spec. + canon. |
| TIDDGIFEV | HSPA1A;SNHG32 | 9 | HLA-C*05:01 | 4.6350 | 4 | 4 | Cell-spec. + canon. |
| RQMKNIVHNY | EPN1 | 10 | HLA-A*30:02 | 4.3566 | 4 | 4 | Cell-spec. + canon. |
| KVKEDVVSY | MYO7A | 9 | HLA-A*30:02 | 3.5080 | 4 | 4 | Cell-spec. + canon. |
| VGDIPGVRF | RPS23 | 9 | HLA-C*05:01 | 3.0428 | 4 | 4 | Cell-spec. + canon. |
| KQYPPNTSKVY | NAMPT | 11 | HLA-A*30:02 | 3.1367 | 4 | 3 | Cell-spec. + canon. |
| SASPIFTHVY | CWC22 | 10 | HLA-A*30:02 | 2.7373 | 4 | 3 | Cell-spec. + canon. |

**Supplementary Table 2.**
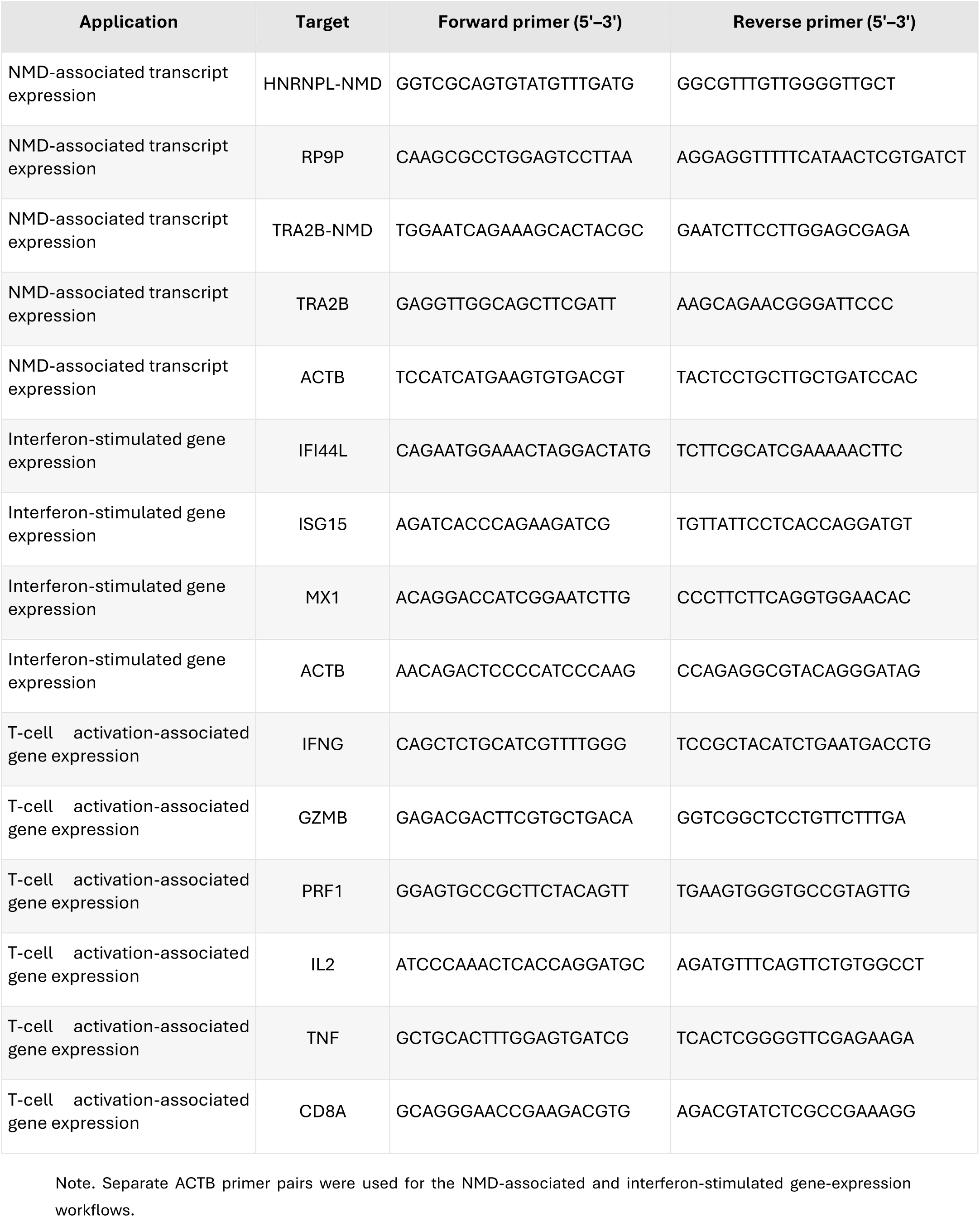
Primer sequences used for RTqPCR.

**Supplementary Table 3.**
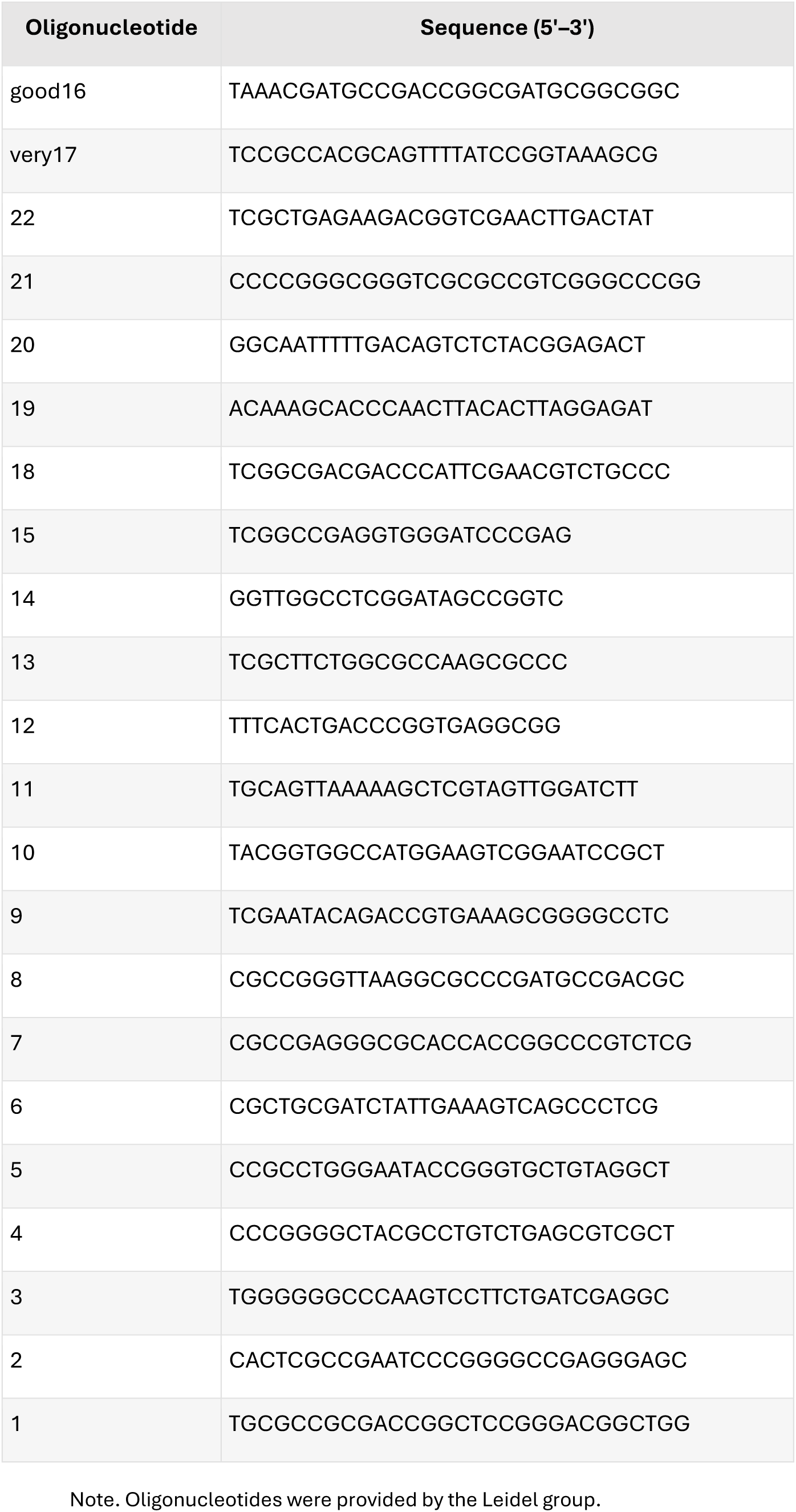
Oligonucleotides used for rRNA depletion.

